# Shielding working memory from distraction is more effortful than flexible updating

**DOI:** 10.1101/743120

**Authors:** Danae Papadopetraki, Monja I. Froböse, Andrew Westbrook, Bram B. Zandbelt, Roshan Cools

## Abstract

Exerting cognitive control is known to carry a subjective effort cost and people are generally biased to avoid it. Recent theorizing suggests that the cost of cognitive effort serves as a motivational signal to bias people away from excessive focusing and towards more cognitive flexibility. We asked whether the effort cost of stable distractor resistance is higher than that of flexible updating of working memory representations. We tested this prediction by using (i) a delayed response paradigm in which we manipulate demands for distractor resistance and flexible updating, as well as (ii) a subsequent cognitive effort discounting paradigm that allows us to quantify subjective effort costs. We demonstrate, in two different samples (28 and 62 participants) that participants discount tasks both high in distractor resistance and flexible updating when comparing with taking a break. As predicted, when directly contrasting distractor resistance and flexible updating the subjective cost of performing a task requiring distractor resistance is higher than that requiring flexible updating.

## Introduction

Cognitive control often refers to the set of mechanisms required to focus on and pursue a goal, especially in the face of distraction, temptation or conflict. Succeeding to exert cognitive control and focusing on the task at hand is highly valued in our industrialized society, as it allows us to complete our tasks and achieve our long-term goals. Despite its importance, failures of cognitive control are very common. Procrastinating, failing to meet deadlines, and performance decrements after fatigue are familiar examples.

Why do people fail to exert cognitive control? Control-demanding tasks carry an effort cost, making people perceive them as aversive^1,2^ and biased to avoid them^3^, even if such avoidance implies forgoing rewards^4,5^. The mechanisms underlying these cognitive effort costs remain elusive. While poor performance on cognitive control tasks has often been explained as a limitation in cognitive capacity, more recent accounts shift the focus from capacity to motivation^6^. These accounts are supported by experiments that show that performance decrements (caused by effort) can be overcome by increases in incentive motivation, for example as a function of monetary rewards^7^. According to resource allocation accounts, the subjective cost of cognitive effort might represent a motivational signal to remain open to alternative opportunities, thus promoting flexibility even at the expense of reduced engagement in a current ongoing task^3,8–10^. As our attentional resources are limited^11,12^, focusing on a given task means that we have to give up on other tasks that require the same set of mechanisms, thus incurring an opportunity cost^9^. Hence, failures of cognitive control can be viewed as stemming not just from failures in *implementation*, but also from a *choice* to pursue alternative tasks that may be more rewarding.

Such a motivational mechanism would be adaptive, given that our constantly changing environment requires a dynamic balance between the cognitive states of focus and flexibility^10,13^. Focusing is crucial for completing our goals, but flexibility is essential when goals change. Flexibility also allows us to explore alternative ways to solve a problem and come up with new ideas, i.e. to be innovative and creative^14,15^.

According to current theorizing^16^, the stability/flexibility tradeoff in working memory is moderated by the strength of current task representations. Strong representations facilitate focusing on a current task-set at the cost of reduced flexibility, for example when task-switching. Weak representations, in contrast, allow flexible adaptation but reduce focused intensity.

How do we decide when to be focused and when to relax constraints in order to be flexible? We have previously argued that we arbitrate between a focused (closed) state versus a flexible (open) one, based on a cost-benefit analysis in which the benefit of cognitive effort corresponds to increased focus and is weighted against its (e.g. opportunity) cost, corresponding to reduced flexibility^13^. We thus reasoned that the cost of distractor resistance is higher than that of flexible updating. Here, we investigate this hypothesis by using a novel version of the cognitive effort discounting paradigm (COGED)^5^ that allowed us to measure the cost that people assign to performing tasks requiring the stable distractor resistance or flexible updating of working memory representations. The process impurity of the classic n-back task^17,18^, used in the original COGED, does not allow this issue to be addressed, because it simultaneously requires letting go of previous representations as well as maintenance and updating. The present design allowed us to separately quantify the subjective cost of a task requiring distractor resistance and that of a task requiring flexible updating. We obtained two independent datasets to replicate, and robustly establish the predicted differences between the costs of focus and flexibility in working memory.

As in the case of the original COGED paradigm, our paradigm consists of two stages. In the first stage, subjects perform variants of a well-established colour wheel working memory task^19^. Participants experience different demands (set sizes 1 to 4) of the two conditions of the task. One condition requires flexible updating; the other condition requires focused distractor resistance (stability). In the second stage, participants make a series of choices between repeating one of the working memory conditions in return for monetary rewards. Some trials require choices between either one of the (ignore or update) task conditions versus taking a break. Inspired by the opportunity cost framework^9^, we opted to offer a time-matched break instead of a lower effort task to incur maximum opportunity costs. Other trials require direct comparisons between the two (ignore versus update) task conditions. In line with our prediction, we show that the cost of distractor resistance is higher than the cost of flexible updating.

## Results

### Working memory task performance

A modified color wheel task was employed (Figure 1A, see Methods section for more details), during which participants were exposed to conditions requiring either distractor resistance (i.e. ignore condition) or flexible updating (i.e. update condition). Every trial of the paradigm consisted of three phases that were separated by two delay periods. In the first phase (*encoding*), participants saw coloured squares, which they always had to memorize. Then after a delay of two seconds, participants saw new colours in the same square locations (*interference phase*). In the ignore condition, participants were instructed to maintain in their memory the colors from the encoding phase and not be distracted by the new interfering colors. In the update condition, participants had to let go of their initial representations and update the new stimuli into their working memory. We manipulated the working memory demand by varying the number of stimuli that needed to be remembered. During the response phase, participants had to match the color of one of the relevant squares by clicking with the mouse on a color wheel.

**Figure 1.**
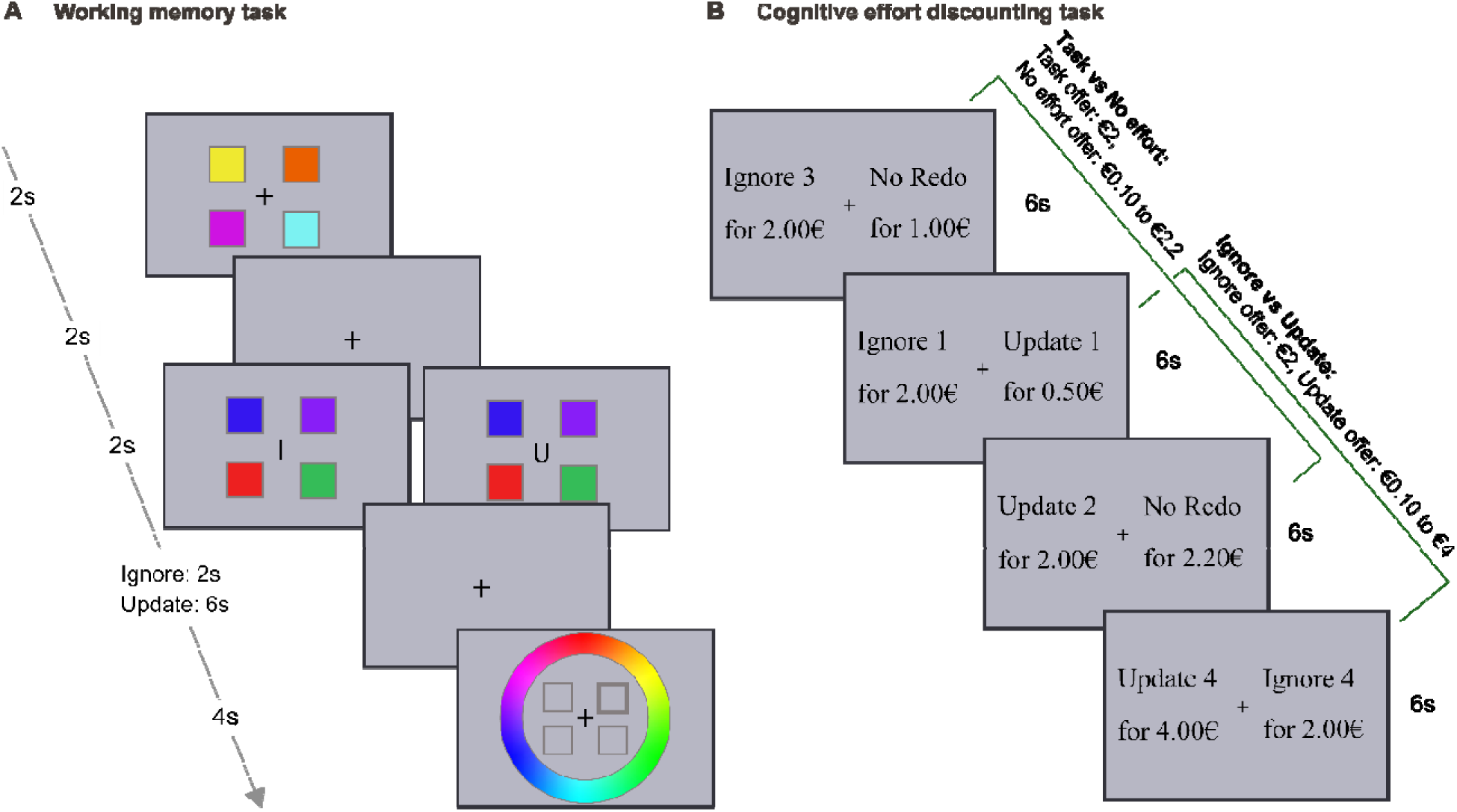
**A** An illustration of the color wheel working memory task. Every trial of the task consists of three phases. In the encoding phase (2 s), participants need to memorize colored squares. After a delay of 2 s, during the interference phase (2 s) a letter indicates if it is an ignore (I for ignore) or an updating (U for update) trial In ignore trials, participants need to maintain colors from the encoding phase and ignore intervening stimuli. In update trials, participants have to let go of their previous representations and update into their memory the stimuli from the interference phase. Another delay separates interference from the response phase. This delay is 2 s for ignore and 6 s for update trials to match the time that the target stimuli are maintained between conditions. During response phase participants see a color wheel and black frames of the same squares; they have 4 s to click on the target color for the highlighted square. Demand is manipulated by varying the number of squares from one to four. The example displayed here is of the highest demand. **B** Example trials of the COGED task. Participants perform two versions of choices. In the “task vs no effort” version, participants have to choose between repeating a level of ignore or update and not repeating the color wheel task at all (“No Redo”). The task option offer remains fixed at €2 and the no effort “No Redo” option varies from €0.1 to €2.2. In the “Ignore vs Update” trials participants must choose between repeating either the ignore or update condition of the same demand. Ignore offers are always fixed at €2 and update offers vary from €0.1 to €4. Trial duration is 6 s. The trials are intermixed.

#### Accuracy

Performance on the working memory task was sensitive to the demand (i.e. set size) manipulation and, in line with earlier studies contrasting ignore and update trials, participants performed more poorly in the ignore compared with the update condition^20,21^ (Figure 2A&B; Supplemental Table 1 for descriptive statistics; Supplemental Figure 1 and 2 for precision indices and signed deviance respectively). This observation was supported by Bayesian model comparison (Table 1), showing strongest support for the model including set size and condition in both studies (BF_10_ = 24876 and BF_10_ = 5.5e⍰+12, respectively). The runner-up model was the one including both main effects and their interaction, which was ∼3.2 and ∼2 times less likely than the model with the main effects only for experiment 1 and 2 respectively. The effects analysis confirmed the conclusion based on model comparison, showing that accuracy decreased with increasing set size (Experiment 1: F_1.52,39.6_= 6.510, p = 0.007, BF_INC_ = 83, Experiment 2: F_1.63,97.7_ = 16.998, p = 2.8e-6, BF_INC_ = 1.6e⍰+10) and that participants performed better on update compared with ignore trials (Experiment 1: F_1,26_ = 11.068, p = 0.003, BF_INC_ = 448, Experiment 2: F_1,60_= 24.095, p = 7.4e-6, BF_INC_ = 939) (Table 2). Evidence for an interaction effect was not conclusive (Experiment 1: F_2.14,78_ = 2.205, p = 0.116, BF_INC_ = 1.3, Experiment 2: F_2.32,139.3_= 3.238, p = 0.035, BF_INC_ = 1.9). We conclude that accuracy decreased as a function of set size and, across set sizes, was worse on ignore relative to update trials.

**Figure 2.**
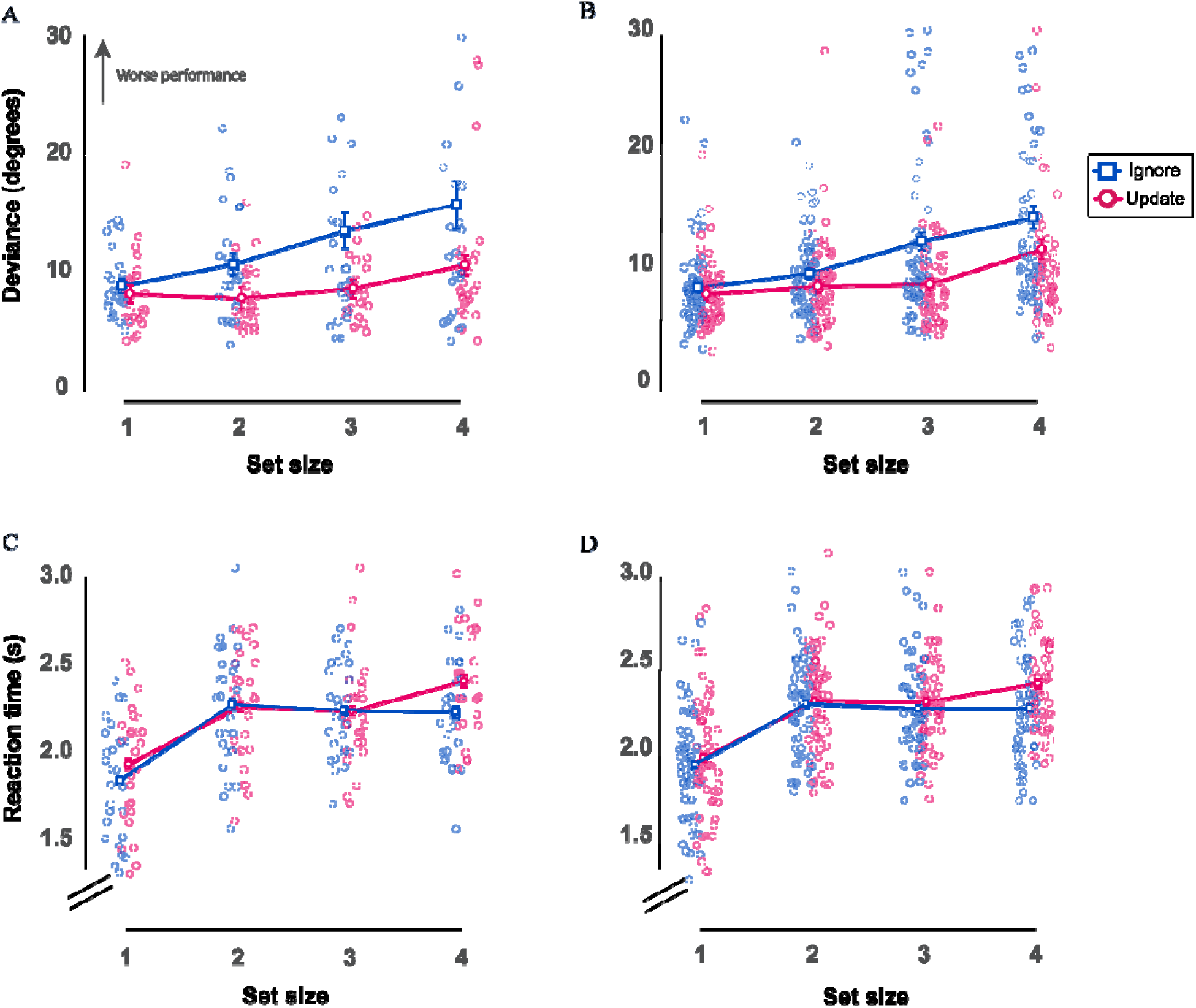
Performance on the color wheel working memory task. **A** Median deviance for experiment 1 (27 participants). **B** Median deviance for replication experiment 2 (61 participants). Deviance in degrees from the correct color is displayed here as a function of set size for ignore and update trials. **C&D** Median reaction times as a function of set size for ignore and update conditions. **C** Experiment 1 (27 participants). **D** Experiment 2 (61 participants). Error bars indicate within-participant SEM^68,69^.

**Table 1.** Model comparison relative to the null model for deviance in the working memory task.

| Models | BF<sub>10</sub> |  |
| --- | --- | --- |
|  | Experiment 1 | Experiment 2 |
| Condition+ Set size | 24875.5 | 5.5e<sup>+12</sup> |
| Condition+ Set size+ Condition* Set size | 8023.0 | 2.7e<sup>+12</sup> |
| Set size | 48.3 | 5.8e<sup>+9</sup> |
| Condition | 263.2 | 346.2 |

**Table 2.** Effects analysis for deviance in the working memory task.

| Effects | Experiment 1 |  |  | Experiment 2 |  |  |
| --- | --- | --- | --- | --- | --- | --- |
| | BF<sub>inclusion</sub> | P<sub>value</sub> | $\eta^2$ | BF<sub>inclusion</sub> | P<sub>value</sub> | $\eta^2$ |
| Condition | 448.2 | 0.003 | 0.299 | 952.0 | 7.4e-6 | 0.287 |
| Set size | 83.2 | 0.007 | 0.200 | 1.75e+10 | 2.8e-6 | 0.221 |
| Condition * Set size | 1.3 | 0.116 | 0.078 | 1.953 | 0.035 | 0.051 |

#### Reaction times

Statistical analyses suggest that RTs varied as a function of demand and task condition; participants were responding faster on trials that presented fewer squares (i.e. lower set size) and in the ignore (versus update) condition (Figure 2C&D; Supplemental Table 2 for descriptive statistics). In the first experiment, Bayesian model comparison (Table 3) showed that the best model was the one including condition, set size and the interaction between the two (BF_10_ = 4.8e⍰+31, ∼1.4 times better than the one also including the interaction). Effects analyses (Table 4) confirmed that participants were faster on ignore compared with update trials (F_1,26_= 16.436, p = 4.1e-4, BF_INC_= 44.6), a very strong set size effect (F_3,78_ = 64.739, p = 4.1e-21, BF_INC_ = ∞) and an interaction effect (F2.41,62.7 = 5.643, p = 0.003, BF_INC_ = 26). In the second experiment, the main effects were in the same direction (set size: F_2.4,141_= 90.386, p = 1.6e-28, BF_INC_ = ∞, condition: F_1,60_ = 16.179, p = 1.6e-4, BF_INC_ = 88), but the evidence for an interaction was weaker (F_2.7,165_ = 4.405, p = 0.007, BF_INC_ = 3.8). The model that only involves condition and set size was marginally better than the one including the interaction. Thus, the dependence of the set size effect on task demand is not clear.

**Table 3.** Model comparison relative to the null model for RTs in the working memory task.

| Models | BF<sub>10</sub> |  |
| --- | --- | --- |
|  | Experiment 1 | Experiment 2 |
| Condition+ Set size | 6.7e<sup>+30</sup> | 1.4e<sup>+53</sup> |
| Condition+ Set size+ Condition* Set size | 4.8e<sup>+31</sup> | 1.3e<sup>+53</sup> |
| Set size | 8.2e<sup>+29</sup> | 2.0e<sup>+51</sup> |
| Condition | 0.84 | 3.5 |

**Table 4.**
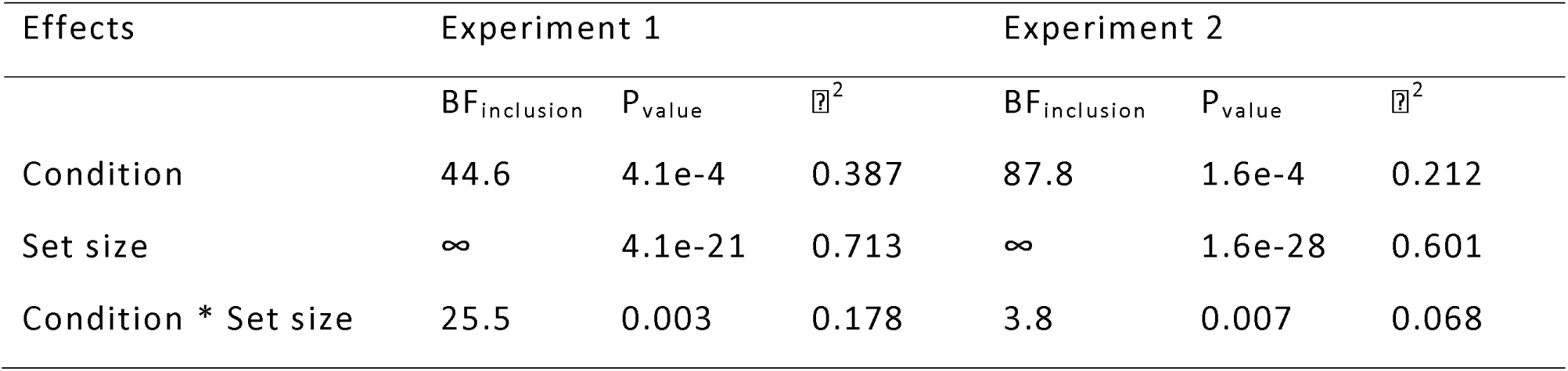
Effects analysis for RTs in the working memory task.

| Effects | Experiment 1 |  |  | Experiment 2 |  |  |
| --- | --- | --- | --- | --- | --- | --- |
| | BF<sub>inclusion</sub> | P<sub>value</sub> | $\eta^2$ | BF<sub>inclusion</sub> | P<sub>value</sub> | $\eta^2$ |
| Condition | 44.6 | 4.1e-4 | 0.387 | 87.8 | 1.6e-4 | 0.212 |
| Set size | ∞ | 4.1e-21 | 0.713 | ∞ | 1.6e-28 | 0.601 |
| Condition * Set size | 25.5 | 0.003 | 0.178 | 3.8 | 0.007 | 0.068 |

### Cognitive effort discounting: Repeat the task or take a break?

Next, we quantified the subjective cost participants assigned to performing the update and ignore trials. The design of this task was inspired by the temporal and cognitive effort discounting literature^5,22^. To assess subjective value, participants made choices about repeating a level of the color wheel task for a monetary reward (effort option) or not repeating it for a usually smaller reward (no effort option) (Figure 1B). We decided to contrast the task offer against a break (instead of a lower load offer) as this reflects real-life choices more closely and incurs higher opportunity costs. The task offer was fixed at €2 and the “no effort” offer varied from €0.1 to €2.2. Every choice was sampled three times to account for response variability. Participants were instructed that after all choices were completed, one of them would be randomly selected and they would repeat a few blocks of that set size and mostly that condition (to reduce predictability). If the “no effort” option was selected, they were instructed that they should remain in the testing room for the same amount of time, but they could use their time as they pleased, e.g. make use of their phone or lab computer. They were also informed that receiving the monetary reward would not be contingent on their performance, as long as they put effort into doing the task.

We computed participants’ indifference points (IPs) to estimate subjective value. Indifference points reflect the monetary amount offered for the presumably less effortful option at which participants are equally likely to choose one or the other, thus the probability of accepting either option would be 0.5. We calculated the probabilities of accepting the presumably easier offer using binomial logistic regression analysis.

The indifference point indexes the degree of discounting of the high-effort offer, where an IP of 2 corresponds to subjective equivalence (given that the offer of the discounted task was always €2, IP = €2 implies that the participant finds the task and the no effort option equally costly). IPs smaller than €2 represent greater discounting (the participant finds the task option to be more costly than the no-redo option) and thus reduced subjective value. Figures 3A&B depict the logistic regression curves of a representative participant for whom it was possible to estimate indifference points (the participant selected both the task and no effort options enough times to fit a logistic regression, see Methods section) for both update (A) and ignore (B) conditions.

**Figure 3.**
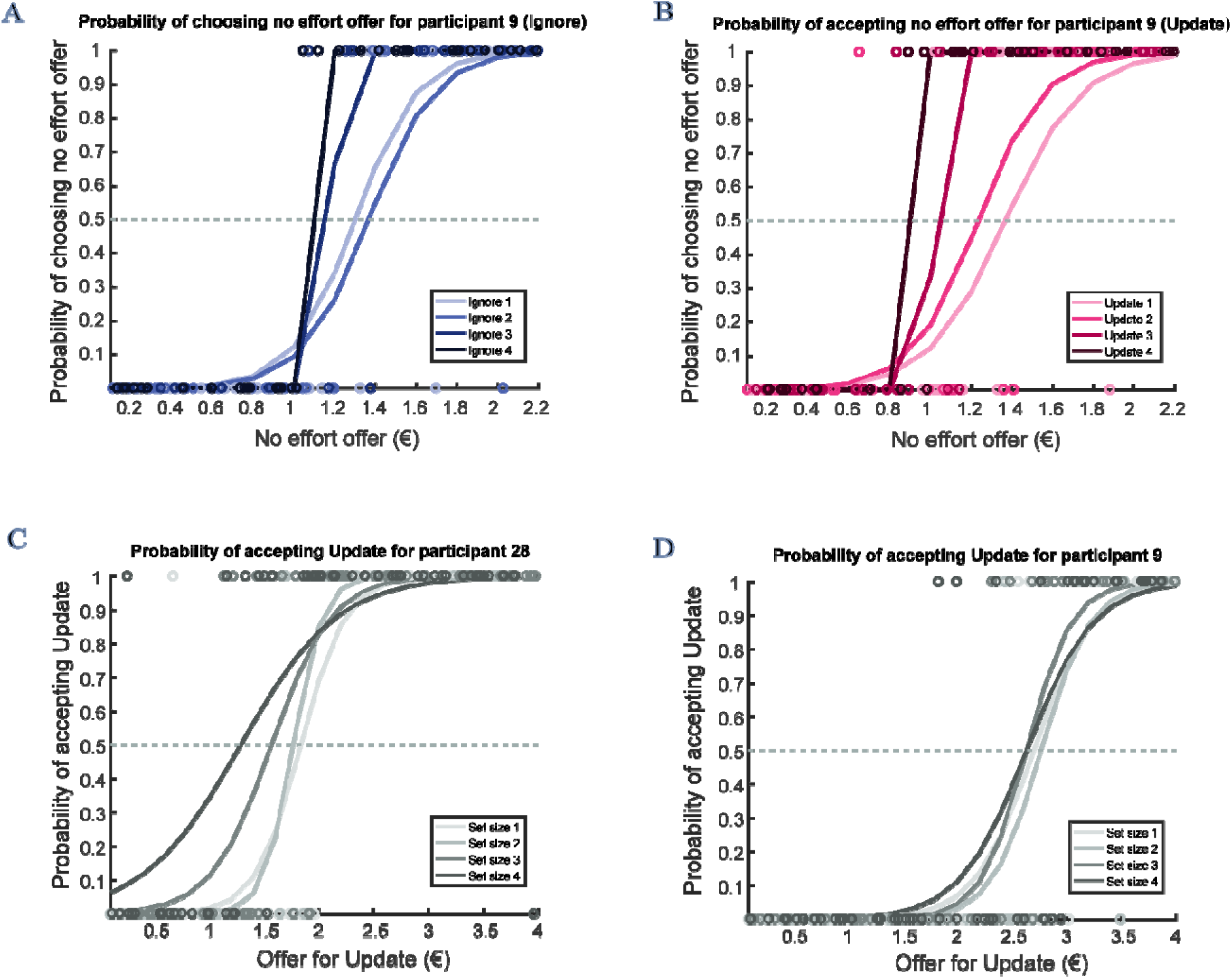
Example participant logistic regression curves. **A, B** Logistic regression curves for “task vs no effort” choices of a representative participant for update (A) and ignore (B) condition. The probability of accepting the “no effort” (i.e. no task) offer (y-axis) is plotted as a function of the amount of money offered for “no effort” (x-axis). Task offer was always €2 for both conditions and all set sizes and the “no effort” offer varied from €0.10 to €2.20. The estimated indifference point is the offer for “no effort” where the probability of choosing to do the task or the “no effort” option is equal (i.e., 0.5). Indifference points decreased with increasing set size. C, D Example logistic regression curves for “ignore vs update” indifference points. The probability of choosing the update offer is shown to vary as a function of the amount of money offered for update. Ignore offer was always €2 for all set sizes, while the update offer varied from €0.10 to €4. The indifference point is the update offer for which the acceptance probability is 0.5, i.e., subjective equivalence. C: Representative participant who discounted rewards in order to avoid ignore trials (preference for update). D: Example participant who discounted rewards in order to avoid the more demanding levels of update trials (preference for ignore).

Next, we analyzed IPs using Bayesian and classical 2×4 repeated measures ANOVAs to assess whether the subjective value of an offer decreased with demand. Indifference points are displayed as a function of set size and experiment in Figure 4A&B (Supplemental Table 3 for descriptive statistics). Overall, the results show that participants found higher demands of the task more costly as the subjective value decreased with increasing task difficulty (i.e. set size). Moreover, in line with our hypothesis, the subjective cost of performing the ignore condition is higher than that of the update condition (Figure 4A to D). On average, participants found the no effort option less costly than the task option, for both conditions. Analyses of data from Experiment 1 (Table 5) showed that the winning model, which included set size and condition (BF_10_ = 5006) was four times more likely than the runner-up model which included set size alone (BF_10_ = 1229). Our second experiment replicated this finding, with the same winning model (BF_10_ = 9.7e⍰+19) being ∼19 times more likely than the runner up (Table 5). Individual effects analyses (Table 6) strengthened these model comparison-based inferences: they provide very strong evidence for a set size effect (Experiment 1: F_1.3,31_ = 5.666, p = 0.016, BF_INC_ = 1246, Experiment 2: F_1.57,77_ = 22.230, p = 2.8e-7, BF_INC_ = 6.0e⍰+15), indicating that participants found higher set sizes to be increasingly costly. In Experiment 1, there was anecdotal evidence that the subjective value of the ignore condition was lower than that of the update condition (F_1,23_ = 10.924, p = 0.003, BF_10_ = 3.1). The more powerful replication study showed extreme evidence for a lower subjective value of ignore versus update (F_1,49_ = 18.216, p = 9.0e-5, BF_10_ = 1684), solidifying the conclusion that participants found the ignore condition subjectively more costly than the update condition. Finally, there is limited evidence against an interaction effect (Experiment 1: F_3,69_ = 1.798, p = 0.156, BF_INC_ = 0.2, Experiment 2: F2.66,130 = 2.167, p = 0.102, BF_10_ = 0.5).

**Figure 4.**
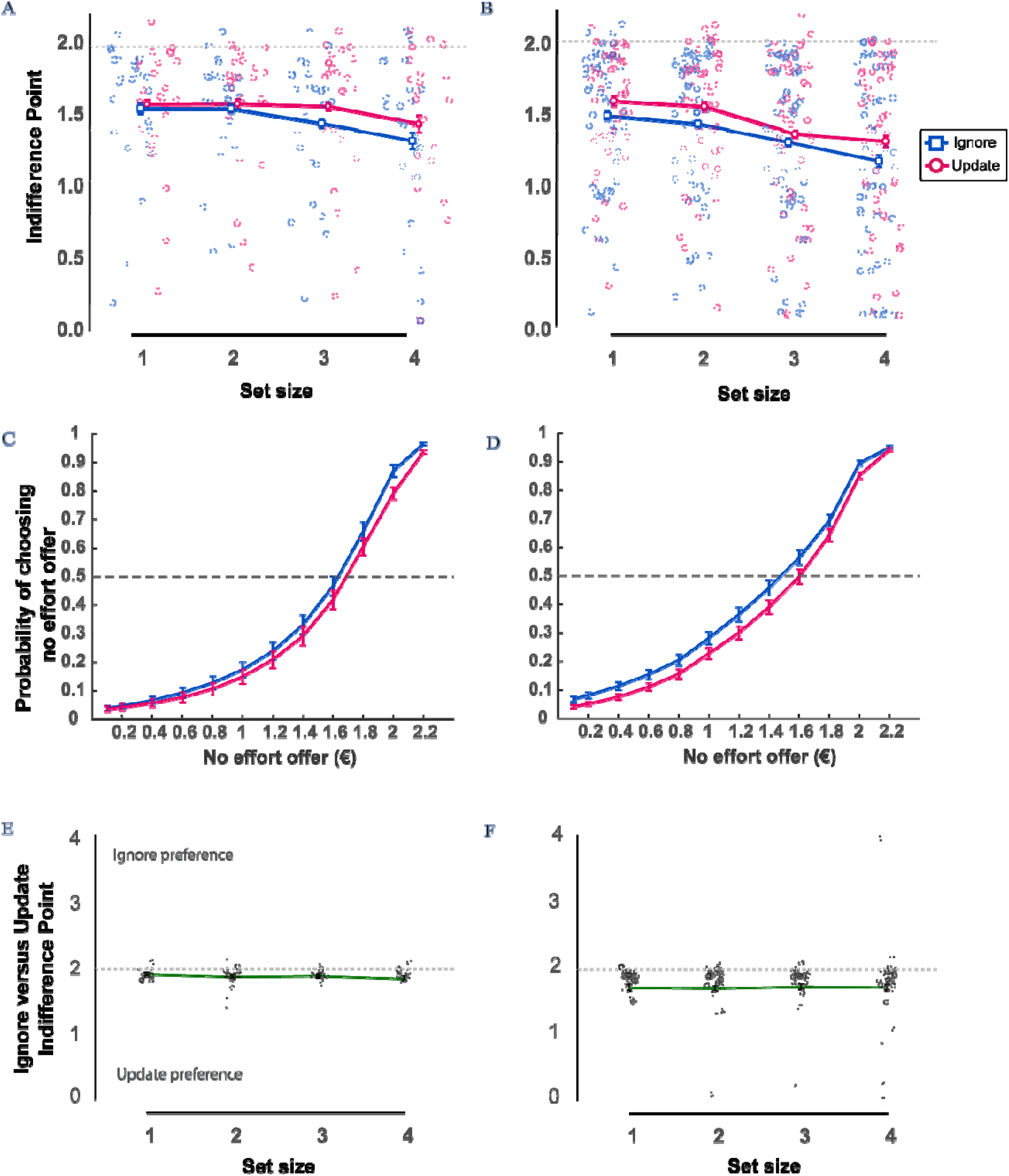
**A, B** “Task vs no effort” indifference points as a function of set size. **A** Experiment 1 (24 participants). **B** Experiment 2 (50 participants). The more the indifference points deviate from 2, the more participants discounted the task option (the task offer was fixed at €2). **C, D** Logistic regression curves for “task vs no effort” choices per condition across set size. The probability of accepting the “no effort” (i.e., no task) offer (y-axis) is plotted as a function of the amount of money offered for “no effort” (x-axis). **C** Experiment 1 (24 participants). **D** Experiment 2 (50 participants). **E, F** Indifference points for “ignore versus update” choices as a function of set size. **E** Data from experiment 1 (26 participants). **F** Data from experiment 2 (58 participants). Indifference points smaller than 2 indicate a preference for update over ignore (offer for ignore was fixed at €2). Error bars indicate within-participant SEM^68,69^.

**Table 5.**
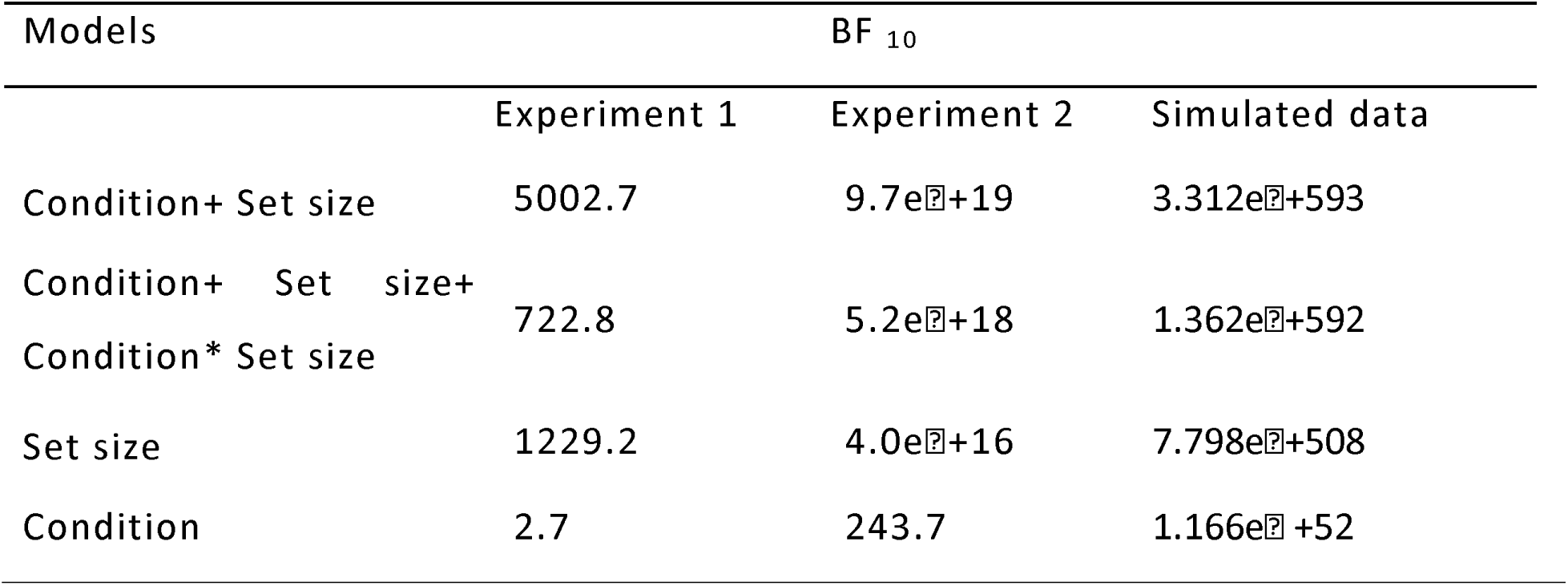
Model Comparison relative to the null model for “task vs no effort” indifference points.

| Models | BF<sub>10</sub> |  |  |
| --- | --- | --- | --- |
|  | Experiment 1 | Experiment 2 | Simulated data |
| Condition+ Set size | 5002.7 | 9.7e<sup>19</sup> | 3.312e<sup>+593</sup> |
| Condition+ Set size+ Condition* Set size | 722.8 | 5.2e<sup>+18</sup> | 1.362e<sup>+592</sup> |
| Set size | 1229.2 | 4.0e<sup>+16</sup> | 7.798e<sup>+508</sup> |
| Condition | 2.7 | 243.7 | 1.166e<sup>+52</sup> |

**Table 6.**
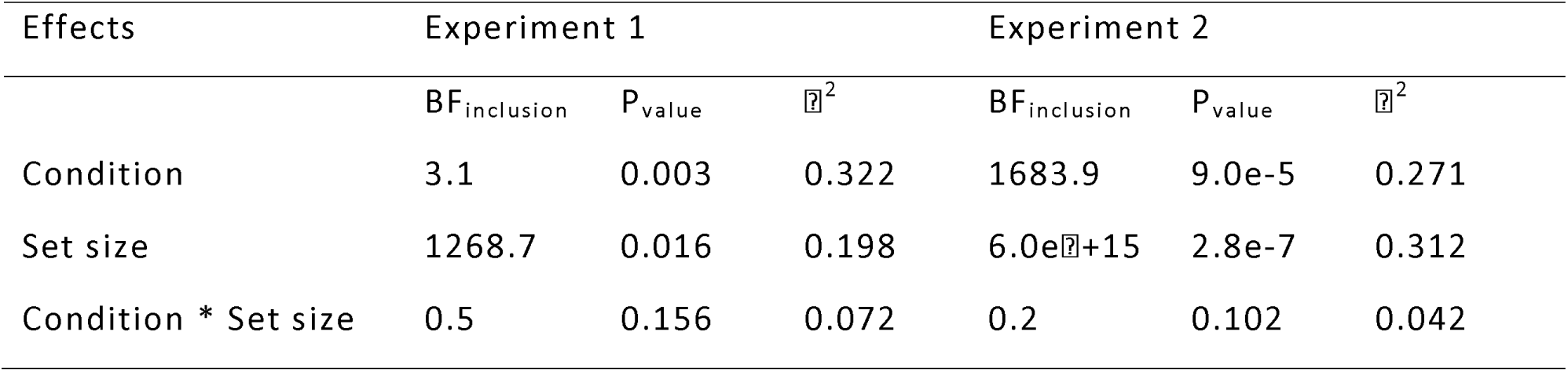
Effects analysis for “task vs no effort” indifference points.

### Cognitive effort discounting: To Ignore or to Update?

Next, we assessed choices that involved direct comparison between performing the ignore and the update trials. For each ignore versus update comparison the set size was identical for both conditions (e.g. ignore 3 vs update 3). The offer for ignore was fixed at €2 and the offer for update varied from €0.1 to €4. Accordingly, an IP < 2 indicates a preference for (increased subjective value of) update vs ignore, while an IP > 2 represents a preference for ignore vs update (see Methods section for more details). Figures 3C&D depict logistic regression curves of two example participants, one preferring the update condition and exhibiting an effect of set size (left panel) and the other preferring the ignore condition and not exhibiting an effect of set size (right panel).

Descriptive statistics are presented in Supplemental Table 4 and one-sample t-test output in Table 7. In Figures 4E&F, we report the average indifference points per set size. In accordance with our second hypothesis, the overall average subjective value of ignore versus update choices was less than 2 (1.88), indicating a preference for update over ignore. The support in the data for this hypothesis is ∼4.8 times higher than the null (T-test (IP<2) t_25_ = −2.440, p = 0.011, BF_-0_ = 4.8). In Experiment 2, the average subjective value was 1.73 and a preference for update over ignore was ∼65 times more supported by the data than no preference (T-test (IP<2): t_57_ = −3.535, p = 4.1e-6, BF_-0_ = 65). The output of the one-way repeated-measures ANOVA shows very strong evidence for the data under the null hypothesis that subjective value is not influenced by set size (Experiment 1: F_1.8,45_ = 0.961 p = 0.382, BF_10_ = 0.149, Experiment 2: F_1.2,69_ = 0.069, p = 0.840, BF_10_ = 0.023), although supplementary analyses revealed that this was due to diametrically opposite effects of set size in participants who preferred ignore and those that preferred update (Supplementary Note 1). Our results increase confidence in our second hypothesis that participants discount rewards in order to repeat flexible updating trials over distractor resistance.

**Table 7.** One-sample t-test results that "Ignore vs Update" IPs are smaller than 2.

| Set size | Experiment 1 | Experiment 2 |
| --- | --- | --- |

|  | BF<sub>o</sub> | P<sub>value</sub> | Cohen's d | BF<sub>o</sub> | P<sub>value</sub> | Cohen's d |
| --- | --- | --- | --- | --- | --- | --- |
| Across | 4.8 | 0.011 | -0.479 | 64.9 | 4.1e-4 | -0.464 |
| 1 | 2.4 | 0.026 | -0.339 | 6698.0 | 2.6e-6 | -0.660 |
| 2 | 4.3 | 0.013 | -0.467 | 1105.0 | 1.8e-5 | -0.588 |
| 3 | 4.2 | 0.013 | -0.464 | 13.2 | 0.002 | -0.508 |
| 4 | 3.0 | 0.020 | -0.426 | 10.9 | 0.003 | -0.488 |

#### Controlling for performance

Having established that ignore is both more difficult and perceived as more costly for most participants, control analyses confirmed that the variability in choice preferences does not merely reflect variability in task performance. Plotting deviance for ignore versus update against preference for ignore versus update reveals little correlation (r = −0.079, BF_10_= 0.181, p = 0.503, Figure 5 & Supplemental Figure 3). We also assessed a relationship between preference and performance using mixed effects logistic regression (see methods). As the design of the experiments is identical, we pooled participants from both studies for these analyses (see separate results in supplementary Note 2). Two models were compared: one including both deviance and condition as predictors of choice and one without condition (see Methods). In the full model, we regressed choice on fixed effects of set size, condition, deviance and no effort monetary reward and added per participant random intercept and by-participant random slopes for the effects of set size, no effort monetary reward and deviance. The reduced model was identical excluding condition. Despite adding deviance to the model, the model including condition was a better fit to the data than the model without condition (reduced model: BIC: 13974, AIC: 13860; full model: BIC: 13900, AIC: 13778, χ^2^ = 84.2, p(pr>Chisq) <2.2e-16) (see Supplementary Note 3 for similar results from ‘ignore vs update’ choices). Additionally, in a model with the same fixed effects as the full model and by-participant random slopes for deviance and condition, the effect of condition was still present (z = −4.5, p = 5.85e-6). Note that the same holds for the effect of set size. We regressed choice on fixed factors condition, set size, deviance and monetary reward for ‘no effort’ and added a random intercept per participant with random slopes for condition, deviance and monetary reward. Model fit was better when including set size, despite adding deviance (model without set size: BIC: 14480, AIC: 14366; full model: BIC: 14153, AIC: 14031, χ^2^ = 337.1, p(pr>Chisq) < 2.2e-16. The above suggest that variability in performance does not explain away differences in preference for update versus ignore or the set size effect.

**Figure 5.**
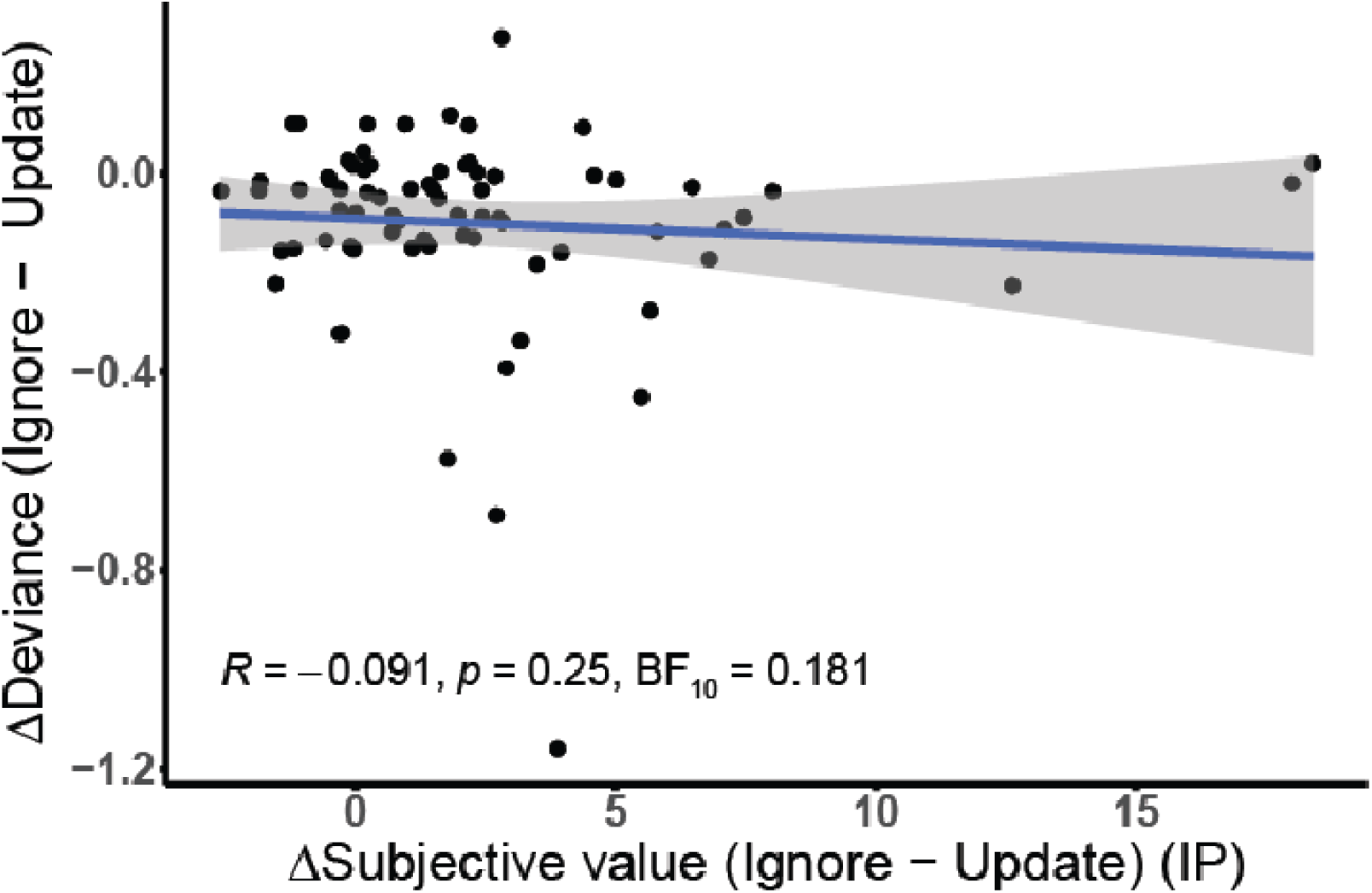
Correlation between condition differences in performance and preference (IP). Ignore minus update performance (median deviance) does not covary with ignore minus update IP. Depicted data pooled from both Experiments (74 participants).

#### Modeling the discounting curve

In addition to our model-free analyses of participant choices, we also pursued an exploratory model-based approach. As there is no definitive consensus on which model best describes the shape of the effort discounting curve, we compared model fits of four models that have been used previously^23–26^ (see methods). The data were best described by the parabolic model including an effort bias parameter^27^ both for ignore and update (Table 8; Supplemental Figure 6). We explored whether the steepness of the discounting curve indexed by parameter k differs between ignore and update conditions. Mirroring our model free results, the k parameter was higher for ignore compared with update indicating steeper discounting of the stable version of the task (BF_10_= 36, z_73_ = 2.982, p = 0.001, Rank-Biserial = 0.399) (Supplemental Figure 7A). We found no conclusive evidence for a difference between conditions for the bias parameter (BF_10_ = 1, z_73_ = −1.98, p = 0.024, Rank-Biserial = −0.265) or the inverse temperature of the softmax function, β, which provides an index of decision noise (BF_10_ = 0.505, t_73_ = 1.350, p = 0.178, Rank-Biserial = 0.181) (Supplemental Figure 7B&C).

**Table 8.** Goodness of fit criteria per condition on “task vs no effort” discounting models.

|  | Ignore |  | Update |  |
| --- | --- | --- | --- | --- |
|  | BIC | AIC | BIC | AIC |
| Linear | 83.2 | 77.3 | 85.6 | 79.7 |
| Parabolic | 92.6 | 86.7 | 93.1 | 87.2 |
| Hyperbolic | 79.5 | 73.5 | 81.6 | 75.6 |
| Exponential | 80.1 | 74.2 | 82.6 | 76.7 |
| Linear + bias | 78.1 | 69.2 | 81.3 | 72.5 |
| Parabolic + bias | 77.9 | 69.0 | 81.0 | 72.1 |
| Hyperbolic + bias | 79.8 | 70.9 | 82.5 | 73.7 |
| Exponential + bias | 78.7 | 69.9 | 81.8 | 72.9 |

#### Does better distractor resistance relate to reduced effort costs?

As an additional exploration, we wondered whether the subjective cost of cognitive effort biases participants towards flexible updating over distractor resistance of working memory representations. Based on a normative computational model of cognitive effort costs, it was proposed that a bias against stable working memory representations is useful for avoiding high switch costs, albeit with greater distractibility^16^. Here, we asked whether those participants who are more sensitive to cognitive effort costs in general (i.e. prefer break over task across conditions)also perform relatively more poorly on stable ignore versus flexible update trials. For this between-subject correlation, we pooled participants across both experiments 1 and 2. We find evidence for this correlation when we focus on the highest effort load (set size 1: Kendall’s tau: −0.005, BF_10_ = 0.15, p = 0.952 ; set size 2: Kendall’s tau: 0.053, BF_10_ = 0.19, p = 0.505 ; set size 3: Kendall’s tau: −0.171, BF_10_ = 1.48, p = 0.031; set size 4: Kendall’s tau: −0.205, BF_10_ = 4.12, p = 0.009)) (Fig. 6). This suggests that people who are worse at ignore compared with update tend to discount high effort tasks more. When examining the correlation of performance and subjective value separately for the ignore and update conditions, we see that the correlation is mainly driven by ignore (set size 1: Kendall’s tau: −0.07, BF_10_ = 0.21, p = 0.409; set size 2: Kendall’s tau: −0.021, BF_10_ = 0.16, p = 0.790; set size 3: Kendall’s tau: −0.228, BF_10_ = 8.75, p = 0.004; set size 4: Kendall’s tau: −0.200, BF_10_ = 6.95, p = 0.012) and not update (set size 1: Kendall’s tau: −0.05, BF_10_ = 0.19, p = 0.499; set size 2: Kendall’s tau: −0.13, BF_10_ = 0.107, p = 0.409; set size 3: Kendall’s tau: −0.141, BF_10_ = 0.72, p = 0.075; set size 4: Kendall’s tau: −0.065, BF_10_ = 0.21, p = 0.414). These results generally concur with the hypothesis that individuals whose preferences are more sensitive to cognitive effort costs, also show greater distractibility (i.e. they are relatively worse at distractor resistance) of working memory representations.

**Figure 6.**
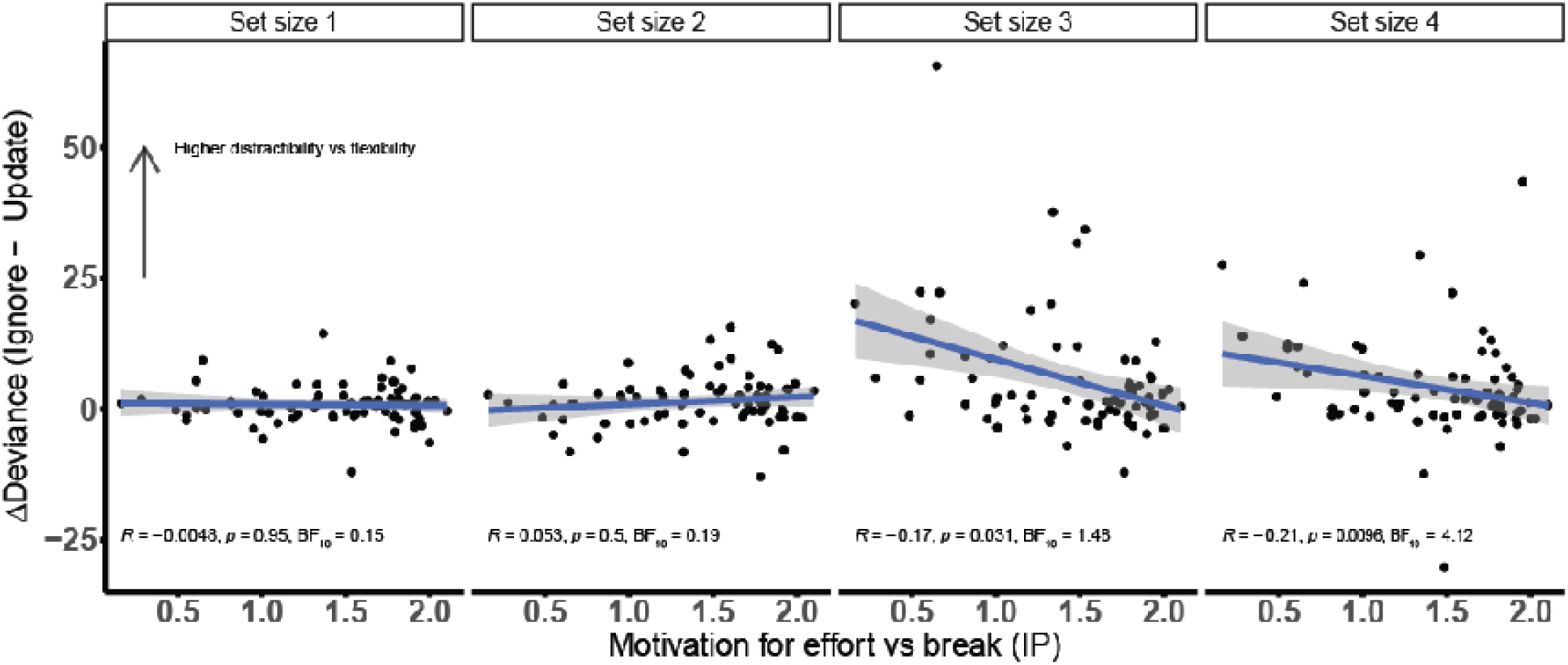
People who show better performance (lower deviance) on the hardest levels of ignore compared with update tend to show higher subjective value for the cognitive task irrespective of condition type.

## Discussion

In this project, we set out to quantify the subjective value of distractor resistance versus flexible updating of working memory representations. We asked not only whether working memory processes are associated with higher subjective costs when demand increases, but also whether tasks requiring flexible updating carry a lower subjective cost than do tasks requiring distractor resistance. In keeping with prior work^4,5,23,28^, we demonstrate highly robust and monotonic discounting of delayed response task value with parametrically increasing working memory load (i.e. set size). Most critically, the results provide strong evidence that the ignore version of the task with high stability demands is more costly than is the update version of the task with high flexibility demands: participants are willing to forgo higher monetary offers in order to avoid repeating performing ignore compared with update trials. This finding is evident both indirectly when participants had to choose between the task and a break, and also directly when they had to choose between ignore and update. This result was replicated in the second independent sample and concurs with our primary prediction that the cognitive effort cost of distractor resistance is higher than that of flexible updating.

The finding that the effort cost of ignoring is larger than that of updating might at first sight seem surprising. First, there is prior evidence for considerable unity of executive functions from individual differences studies^29^. Second, many studies have shown that tasks with high demands for cognitive flexibility, like task switching and set-shifting, are accompanied by robust (residual) costs^30–32^. Additionally, a recent study employing n-back and detection tasks showed that updating may incur higher costs than distractor resistance^33^. However, these tasks require robust maintenance as well as updating for successful performance and inference about differential costs for different cognitive operations depended on model-derived analyses. Our task was specifically designed to address this question by comparing two well-matched conditions differing primarily in demands for flexible updating and distractor resistance. Notably, the working memory load or encoding recency of the trial-relevant stimuli are also matched between the ignore and update conditions Interestingly, we get the expected result despite the demands on working memory input gating being twice as high for the update compared with the ignore trials and update trials being longer compared with ignore trials both by design and due to slower RTs in the update condition. Thus, it would be more precise to state that the cost of ignore is higher than the cost of update plus a time cost.^30–33^.

What then is the origin of the increase in cost for distractor resistance versus flexible updating? One possibility is that this effect reflects a difference in opportunity costs. The more costly ignore trials were 4 seconds shorter than the cheaper update trials, and thus opportunity costs are unlikely to map directly to time costs^34^. Instead, we argue that the cost of maintenance reflects an opportunity cost of focusing: the cognitive strategy required for accurate ignore versus update performance differs in the degree to which it allows novel input to impinge on current processing. More generally, it is possible that the brain is more strongly biased against tasks that demand stable focusing compared with flexible opening given that focusing will incur higher opportunity costs across environments.

The observation that the subjective cost of ignore is higher than that of update is in line with the finding that participants perform more poorly on ignore compared with update trials. The performance effect concurs with previous results from studies using an analogous task with ignore and update conditions^20,21,35^. In those prior studies, however, the task-relevant delay between the to-be-remembered items and the probe was shorter in the update than the ignore condition, rendering inference about the cognitive mechanism underlying the performance difference difficult. Here we show that the ignore condition is accompanied by worse performance than the update condition, even if task-relevant delay is matched between conditions.

A key question that is raised by the performance difference between task conditions is whether the condition effect on subjective effort cost reflects differences in the degree of (aversion to) anticipated performance error. We argue, however, that an increase in the anticipated performance error is unlikely to account fully for the increase in subjective effort cost of the ignore versus update condition, for the following three reasons. First, while instructing participants, we highlighted that monetary rewards would not be contingent on performance during the ‘redo session’, so that performance error should not have influenced participants’ choices in our design. Second, in statistical mixed-effects models that took into account accuracy, the effect of condition was still present.

Third, there was no evidence for an association between performance error and measured preference (Figure 5). In future studies, we might consider matching performance between the two conditions or provide “fake” feedback to influence participants’ beliefs about their performance.

Notably, participants responded not only more accurately, but also more slowly on the update than the ignore trials. We are puzzled by this finding, and consider it possible that the slowing reflects a reduction in a nonspecific orienting response to the intervening stimulus, which might have acted as a warning signal. Warning signals are known to induce slowing of responses as a function of foreperiod (delay), which in our case is longer in update trials^36^. This hypothesis is supported indirectly by the observation that reaction times in flexible update conditions in previous studies in which cue delays were matched between conditions were faster than in ignore conditions^20,21^. We also consider an alternative explanation, namely that the effect of condition on reaction times reflects a modulation of a decision threshold rather than of attentional orienting. Thus participants might trade off time for higher accuracy^37^ in the update condition in which the memory is more robustly maintained and such a strategy would be beneficial.

In addition to disentangling the subjective value of distractor resistance and flexible updating, the present results strengthen and extend previous studies on the value of cognitive engagement. First, we confirm that, on average, people are averse to cognitive demand, are ‘cognitive misers’, willing to decline rewards in order to avoid demanding tasks. This strengthens earlier work showing that participants prefer to avoid more demanding N-back tasks^5,38^, detection tasks^23^ and sustained attention tasks^4^. Our results further generalize these conclusions to the most classic of working memory tasks: the delayed response task. We further generalize these findings to choices between cognitive effort and a time-matched break. Offering a break compared with a low-effort task potentially better reflects choices in daily life, allowing for greater ecological validity and higher opportunity costs. A distinct strength of our design is the fact that our implementation of the discounting procedure takes into account the observation that choices are probabilistic^39^. Unlike prior studies on cognitive effort which used staircase procedures sampling every choice option only once^4,5^, we sampled the full discounting curve and every choice option multiple times. Furthermore, unlike prior studies, in which on the first trial a lower monetary offer was made for the low effort option than for the high-effort option, we avoided (potential) anchor effects by presenting offers randomly. Finally, we also offered participants the opportunity to choose the effortful option for less money. As expected, most participants declined this offer, but the subjective value of four participants (total in both samples) was higher than 2 for at least one of the two working memory processes, indicating a preference for repeating the working memory task, suggestive of effort seeking^40^. Similarly, even though the majority of subjects showed a preference for the flexible condition, there was a small subset of subjects preferring the distractor resistance condition, implying the presence of different subtypes.

Modeling the shape of the discounting curve revealed that the parabolic model including a bias parameter provided the best account of our cognitive effort data. Unlike delay discounting, where a convex pattern has consistently described behavior, effort discounting modeling attempts have been less conclusive^25,26,41,42^. Our results converge with the idea that discounting patterns for effort might differ between tasks and individuals^24^. However, we hesitate to draw strong conclusions about this finding because this was an exploratory analysis and our task was not designed to address this question.

In our subsequent exploratory analyses we saw that participants who are better at high-load ignore versus update trials showed smaller effort costs across set sizes and conditions. This analysis was inspired by normative computational neural network modeling work proposing that the subjective cost of cognitive effort is associated with a bias against distractor resistance and towards flexible updating, thus attenuating switch costs^16,43^, but see^44^. Our result is consistent with the prediction that people who are less sensitive to the subjective costs of cognitive effort, engage more deeply and stably with working memory representations and perhaps show higher switch costs. Conversely, those with more sensitivity to subjective costs are less deeply focused (and thus show greater deviance on ignore trials), and perhaps have less difficulty switching representations flexibly. While our result is consistent with this hypothesis, future studies are needed to directly manipulate task demands for maintenance versus updating and test the effects of such fluctuations on subjective effort costs. A key prediction is that participants experience higher subjective costs of control when the environment is more changeable, and when demands for flexible updating are thus higher.

The neural mechanisms underlying the considerable individual variation in subjective cognitive effort costs should be addressed in future work. Past work has shown that administration of the catecholamine reuptake blocker methylphenidate improves distractor resistance at the expense of flexible updating on a task analogous to the one employed here^21^. However we also know that effects of psychostimulants vary greatly with individual baseline measures of striatal dopamine function^45^. How does a preference for ignore versus update relate to baseline measures of dopamine and psychostimulant effects on cognition? Recent demonstrations that striatal dopamine plays a key role in cognitive effort-based decision-making^46–48^ are consistent with studies in physical effort, where dopaminergic medication has been shown to increases selection of high effort/high reward trials in patients with Parkinson’s disease^49^, while dopamine depletion decreases willingness to exert effort in humans and rodents^50,51^. Another potentially relevant neurotransmitter is noradrenaline which is involved in switching modes between task engagement (focusing) and disengagement (distractibility)^52,53^. Indeed, recent evidence indicates that amphetamine and methylphenidate, both altering catecholamine transmission, modify cognitive demand avoidance in rodents^54^ and humans^46,47,55^ respectively. Together, these findings suggest that catecholamines contribute to valuation of cognitive effort^46,47,55^ and we believe our paradigm is suited to aid uncovering such effects in future research.

In conclusion, this study provides new insights to the novel and growing fields of cognitive effort discounting and value-based decision-making. Specifically, we provide strong evidence that distractor resistance is perceived as relatively costlier than flexible updating, thus contributing considerably to our understanding of the origins of cognitive effort costs.

## Methods

### Participants

For Experiment 1, 32 participants (22 women), aged between 18-29 years old were tested in total. Participants had normal or corrected-to-normal vision. Colorblind participants were excluded. Four data sets were lost during data transfer, so we ended up with 28 data sets (20 women, 18-33 years old, mean: 24). For Experiment 2, we sought to replicate the finding that update is more costly than ignore. We performed a sequential sampling power calculation using the BFDA^56^ package in R. We set the minimum sample size to 60^57^, the maximum to 100 and set the boundary of sampling at a Bayes Factor of 10 for either the null or the alternative hypothesis given the effect size estimated from Experiment 1. We collected 62 data sets (37 women, 20-44 years old, mean: 25.6, standard deviation: 4.3), at which sample size the boundary was already reached. The study was approved by the local ethics committee (CMO region Arnhem/Nijmegen, The Netherlands, CMO2001/095) and all participants provided written informed consent, according to the declaration of Helsinki.

### Exclusion criteria

We excluded participants based on four rules: 1) Failing to pass the color sensitivity test twice. 2) Striking evidence that they did not understand or will to perform the tasks. 3) Mean deviation exceeding 3 standard deviations from average for at least one of our main conditions (across demand) of the working memory task. 4) Failing to estimate reliable indifference points for at least one condition (across demand levels) of the effort discounting tasks.

Based on our criteria, one outlier was excluded from performance analysis of Experiment 1 for deviating more than 3 standard deviations from the mean for ignore (∼3SD) and one from Experiment 2 for deviating more than 3 standard deviations from the mean of both conditions (∼5.4SD from ignore and ∼6.6SD from update mean). Four people were excluded from the analysis of task vs no effort indifference points in Experiment 1 and twelve in Experiment 2. In Experiment 1, all four were excluded because we could not estimate indifference points for at least one of the two conditions (ignore/update). Among the four that were excluded, one always chose the no effort option, one of them always chose the task option and one of them always chose no effort for update trials and task for ignore trials. In Experiment 2, eleven participants were excluded due to inadequate response variability and one because he was not performing the task. Out of the eleven whose IPs we could not estimate, one almost always chose the task option and the rest always preferred the no effort option. The other participant always responded using one of the two response buttons. This is a clear indication that he was not trying to perform the task because easy and hard offer presentation was counterbalanced across response buttons. We excluded two participants from the analysis of “ignore vs update” indifference points in Experiment 1 analysis; one because we could not estimate any indifference points (always chose ignore) and another because they deviated more than 3 standard deviations from the mean. Four participants were excluded in the replication for the same analysis. One always chose ignore, two always chose update and one did not do the task (see above). The participant who did not do the task was the only participant excluded in the mixed models analyses.

### Task design

All paradigms were entirely programmed in MATLAB (Mathworks, Natick, MA, USA)(release 2013a) using the Psychophysics Toolbox extension^58^ (version 3.0.12) on a Windows 7 operating system. The screen resolution was 1920×1080 pixels. The background color for all paradigms was grey (R: 200 G:200 B:200).

The experiment lasted about 130 minutes and consisted of four tasks performed at a computer and questionnaires that participants filled in at the end. The first task (∼7min) was a color sensitivity test aiming to check whether participants were sensitive to the colorful stimuli used in the memory task. Participants then proceeded with the color wheel working memory task to acquire experience with varying demand of the two working memory processes of interest (∼10min practice and 30min task). The third task (∼5min practice and 55min task) was a cognitive effort discounting paradigm that was used to estimate subjective value and address our research questions. The last computer-based task was a redo of the color wheel task (∼10min). Finally, participants filled in some experiment-related questionnaires (∼5min).

#### Color sensitivity task

For our working memory task, we used color stimuli and a color wheel, so it was crucial that our participants’ color vision was not impaired. To test their sensitivity to our manipulation we developed a version of the color wheel task without a working memory component. In this task participants viewed a colored square in the middle of the screen and the same color wheel used in the memory task. Their goal was to click on the color of the wheel that matched the colored square.

The stimuli used for the color sensitivity task were a color wheel, black lines and colored squares. The color wheel was created by 512 successive colored arcs of equal angle (512/360° = 1.42°), each arc carrying a different color. The radius of the wheel was 486 pixels. To form the wheel into a ring, a smaller circle was superimposed, whose radius was ∼362 pixels. The centre of both the wheel and the circle coincided with the centre of the screen. The 512 colors of the color wheel arcs were generated using the hsv MATLAB colormap. The black lines were 0.4° black arcs.

In every trial of this task, participants viewed the color wheel and a colored square in the middle of the screen (Figure 7). They were instructed to look at the color of the square and use the mouse to click on the corresponding shade on the color wheel. To indicate that their response was recorded a black line appeared on the color wheel and successively another black line appeared designating the location of the correct color. Feedback consisted of the actual deviance plus a positive message (‘Good job! You deviated only __ degrees.’) and was provided only when responses deviated less than 10°.

**Figure 7.**
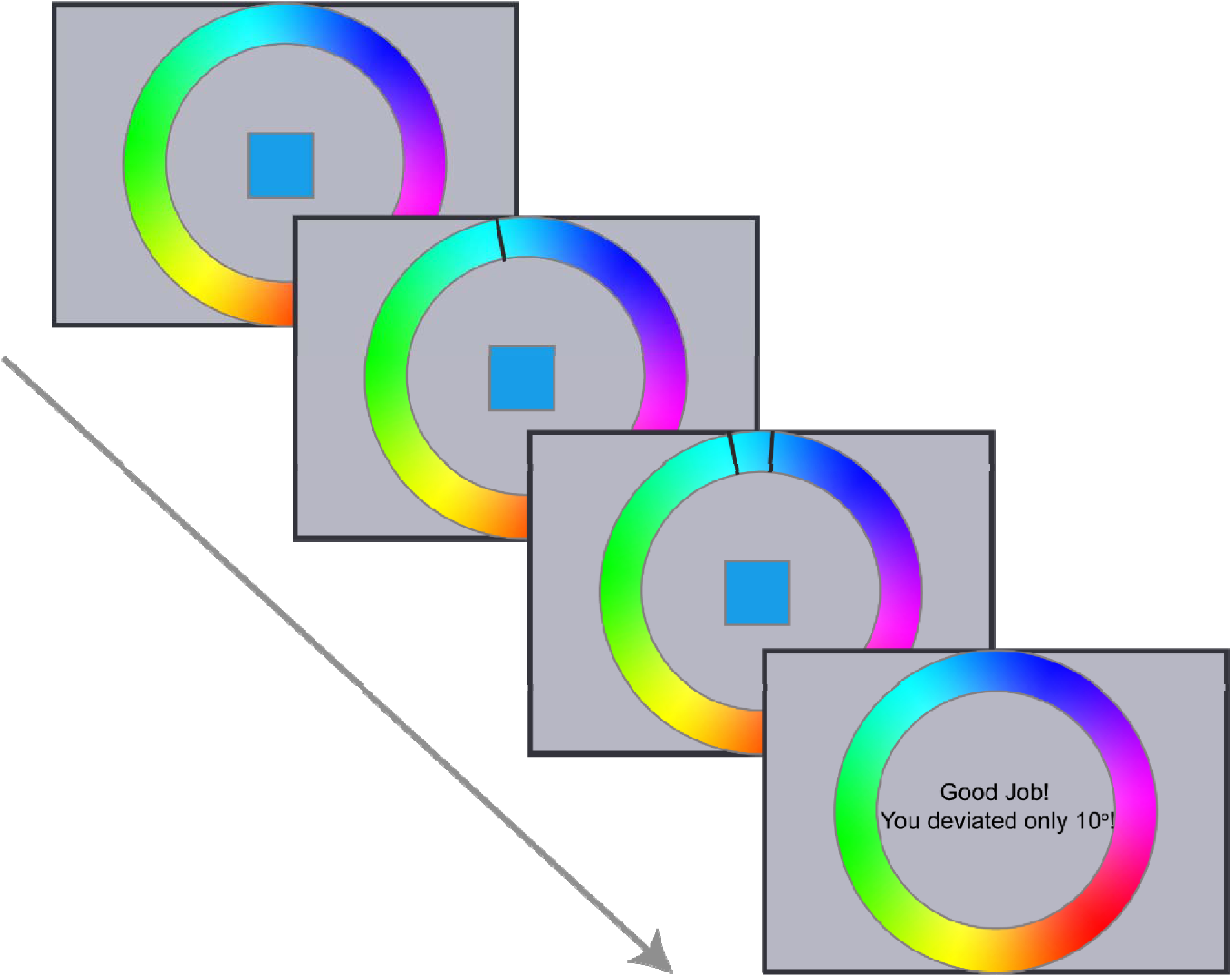
Example trial of the color sensitivity task. Participants view a colored square in the middle of the screen and they have to click with a mouse on the corresponding color on the color wheel. A black line indicates the selected color and a successive line the correct color. If the selected color deviated 10° or less from the correct color they receive feedback that they performed well. The task is self-paced (24 trials total). They successfully complete the task if their average deviance is less than 15°.

To test a representative sample of the color wheel we split the wheel in 12 main arcs. Participants were tested in two different shades from each of the 12 color categories (arcs). So, they performed in total 24 trials of this task. The presentation of the trials as well as the orientation of the color wheel were randomized. The responses were self-paced and total task duration was approximately 7min.

The main dependent variable in this task was deviance in degrees from the correct color. If their average deviance was less than 15° by the end of the task, the experiment continued. Otherwise, they had one more chance to perform the color sensitivity task, but if failed again they would be excluded.

All participants from both experiments completed 24 trials of the color sensitivity task and they all met the criterion (average deviance from correct color below 15 degrees) to continue to the main paradigm. For Experiment 1, the average deviance from the target color was 6.63 degrees (SD = 1.23; median = 4.72, SD = 0.85) and for replication Experiment 2 mean deviance was 6.27 (SD=1.4; median=4.85, SD=1.08). We also reported the median for easy comparison with the color wheel working memory task results.

#### Color wheel working memory task

After successfully completing the colour sensitivity task, participants proceeded with the colour wheel working memory task. In this part, participants experienced varying demands of cognitive stability and cognitive flexibility. This task was based on a short-recall task^19^ and delayed-match-to-sample tasks^20^ that have previously been used to disentangle between the two working memory processes of interest.

The stimuli displayed during this paradigm were a color wheel, colored squares, black frames of squares, a fixation cross, black lines and central letter cues. The color wheel was generated as described in the color sensitivity section. The number of squares varied from one to four and they could be located in four different positions. The centres of the squares formed a rectangle with dimensions 248*384 pixels. Each of the four squares was 100×100 pixels in size. To choose the colors of the squares, we split the color wheel into 12 main arcs of 42 colors each and only used the 15 central colors of each arc. The arcs from which the colors would be sampled per trial were defined manually, but the exact shade (RGB values) was randomly selected. The letter cues were “I” and “U”, colored black and presented at the centre of the screen.

Every trial of the task consisted of three phases separated by two delay periods (Figure 1). During the encoding phase, participants viewed the fixation cross and one to four colored squares for two seconds. The number of squares displayed (set size 1-4) represented the demand of the trial. A delay of two seconds followed, during which only the fixation cross was displayed. Then the interference phase followed. In this phase, participants viewed the same number of squares as during encoding, at the same locations, but with different colors. Instead of a fixation cross, one of the two letter cues was presented during interference in the middle of the screen. The cue indicated the condition of the trial: “I” for ignore trials and “U” for update. The second delay duration depended on trial condition, and was two seconds for ignore and six seconds for update trials. Finally, during the response phase participants saw black frames of the same squares, one of which was highlighted, in addition to the color wheel and the fixation cross. If the participant responded within four seconds, a black line appeared on the color wheel, otherwise, they were instructed to respond faster (‘Please respond faster!’). The total duration of the response phase was five seconds. For the encoding phase, participants were instructed to always memorize the colors and locations of all presented squares. The instructions for the interference phase differed based on the condition as indicated by the letter cue. In ignore trials, participants needed to maintain in their memory the colors from the encoding phase and not be distracted by the new intervening stimuli. In flexible updating trials, participants had to let go of their previous representations and update into their memory the stimuli from the interference phase. Thus, the colors that needed to be remembered for the ignore condition were the ones from the encoding phase, while for the update condition they were the ones from the interference phase. To match the time that the relevant stimuli were maintained in memory for both conditions, the second delay was 4 seconds longer for update trials. Participants were to indicate the color for only the highlighted square. They had to identify the target color on the color wheel and click using a mouse, within four seconds. Only the first response counted. A black line indicated their response. Only during practice trials, a second line appeared at the correct color and positive feedback was displayed if they were performing well (as in color sensitivity section). During the task, no feedback was provided. We instructed participants to fixate in the middle of the screen throughout the task in order to dissuade them from adopting the strategy of closing their eyes during ignore trials in order to avoid being distracted.

Participants first underwent a practice session of 16 trials and then performed two blocks of the task. A block consisted of 64 trials, resulting from repeating each combination of difficulty (four levels: set size 1 – 4) and condition (two levels: ignore and update) eight times. Depending on the difficulty level of the trial, a group of two to eight colors was used to create the trial stimuli, each color coming from one of the 12 arcs. Colors of the same arc never appeared more than once in the same trial. To make sure that ignore and update trials were as similar and counterbalanced as possible, the color stimuli sets displayed and the target colors were the same for both conditions. Because the relevant colors appeared during encoding for Ignore and during interference for update, we made sure that the same group of stimuli also appeared in reverse order between these two phases. So, the same groups of colored squares were presented four times per set size and in total 32 groups of colors were used. To decrease learning effects due to repetition, we split the same stimuli groups between the two blocks. To control for differences between the two hemispheres in representation of color^59^, target locations (left/right) were counterbalanced across conditions. Moreover, the same colors were highlighted for all four set sizes.

#### Cognitive effort discounting task

After participants gained experience with all four difficulty levels of update and ignore conditions of the color wheel working memory task, they proceeded with the third part of the experiment: the effort discounting task (Figure 1B). The aim of this paradigm was to quantify the subjective value that participants assigned to the color wheel task performance. There were two versions of choice trials to address our two research questions. In both versions, two options were accompanied by an amount of money and the options defined what participants would do in the last part of the experiment.

In every trial of the task, participants saw a rectangle containing two options and a fixation cross. The options could be “No Redo” or any set size of ignore or update, for example “Ignore 2”, corresponding to the ignore condition of the task and set size of 2. Below each option, a monetary reward was displayed, for example “for 2€”. Participants could choose the left or right option by pressing 1 or 2 on the keyboard and they had six seconds to respond. When participants made a choice, a black square surrounding the selected offer appeared to indicate that their response was recorded.

At this stage, participants were instructed that there were two more parts in the experiment. In the last part, they would have the opportunity to earn a bonus monetary reward by redoing one to three blocks of the color wheel task. However, the amount of the bonus and the type of trials they would repeat would be based on the choices they made on the choice task. To highlight the importance of every choice, we instructed them that of all the choices they made (of both versions) the computer would select only one randomly and the bonus and redo would be based on that single choice. To minimize effects of error avoidance on choices, we informed participants that accuracy during the redo part would not influence whether they receive the monetary reward, as long as their performance was comparable with the first time that they did the color wheel task (part 2 of experiment). Both the rewards and the redo were real and not hypothetical.

#### Task vs No effort: Choices between working memory task and a break

These trials addressed the first research question: whether the subjective values of ignore and update decrease as a function of task demand. Here, participants had to choose between repeating a level of ignore or update (effort offer) and not redoing the color wheel task at all (no effort offer). If they chose the no effort option (“No Redo”) they were instructed that they would be able to use their time as they pleased (e.g., by using their phones or lab computer) but they would still have to stay in the testing lab so that time spent on the experiment was the same for both options. Otherwise, if the option to repeat the task was selected, the redo trials would consist of mostly (70%) the selected choice condition and level. “Mostly” is important because if they always did the same condition during the redo, they would be able to predict whether they had to update or ignore. We emphasized that they should take their time to respond, consider both the money and their experience while doing the color wheel task as well as the importance of choosing their true preference and not try to please us.

#### Ignore vs Update: Choices between distractor resistance and flexible updating

This COGED trial type aimed to investigate whether ignore is perceived as costlier than flexible updating by directly contrasting them. In these trials, participants had to choose between doing the same level of either ignore or update.

For the ‘task vs no effort’ version of the COGED, the amount offered for the no effort “No Redo” option varied from €0.10 to €2.20 in €0.20 steps (except the first step, which was €0.10), while the task option (effort offer) was always fixed at €2.00. The €2.20 option for “No Redo” was included to identify whether there were participants who strongly preferred performing the task, even if that meant forgoing rewards. As we hypothesized that ignore would be costlier, in the ‘ignore vs update’ version of the task, ignore (hard offer) was kept steady at €2 and update (easy offer) was varying from €0.10 to €4 in €0.20 steps (as above). There were 96 possible pairs for “task vs no effort” choices (12 amounts*2 conditions*4 set sizes) and 84 for “ignore vs update” choices (21 amounts*4 set sizes). Given the evidence that choice is probabilistic rather than deterministic^39^, every pair of options was sampled three times. We decided on three repetitions of the pairs based on a simulation analysis using pilot data (Supplemental Figure 4) in order to optimize the trade-off between indifference point estimation and task duration. Each participant performed three blocks that contained in total 288 trials of “task vs no effort” trials and 252 trials of “ignore vs update”. The trials of the two versions were interleaved (mixed) and randomized within each block. To avoid location effects, we counterbalanced the left-right presentation of the two options. Total task duration was about 55 minutes.

We decided to use fixed sets of offers and not a titrated staircase procedure to estimate subjective value because staircase procedures do not sample the entire logistic regression curve. This COGED version allowed us to sample the logistic regression curves adequately because all participants were faced with the entire range of offer options. The offer range was based on previous discounting studies^5^.

#### Redo

After participants finished three blocks of the discounting choice task, one of their choices was pseudo-randomly selected. Specifically, the computer only sampled from “ignore vs update” choices of level 3 or 4. Participants always did one block of 24 trials of the color wheel task. Two-thirds of these trials were their preferred condition (ignore/update). We decided to never select the no effort option to maintain experimenter credibility, so that participants discussing the task are convinced that the consequences are real. The redo data were not analysed and participants always received the bonus regardless of their performance.

#### Debriefing questions

After the end of the experiments we requested participants to complete questionnaires. We explicitly asked them to report their preference by asking “Which trials did you prefer?”.

### Data analysis

We analysed our data using both frequentist and Bayesian statistics. All statistical analyses were performed using open-source software JASP (version 0.7.5.6)^60,61^ on a Windows 7 operating system.

As skepticism against classical statistical tools increases^62^, we turned to Bayesian statistics^63^. This allowed us to quantify evidence for our hypotheses instead of forcing an all-or-none decision and an arbitrary cut-off of significance. Bayesian statistics can also provide evidence for the null hypothesis (H_0_), thus distinguishing between undiagnostic data (“absence of evidence”) and data supporting H_0_(“evidence of absence”). Another important benefit is that we are able to monitor evidence as data accumulate and we can continue sampling without biasing the result. Due to all the above advantages, we decided that our main conclusions would be drawn based on the Bayesian analyses.

However, frequentist statistics are well-established and widely-acknowledged tools, so more scientists are familiar with their rationale and interpretation. To ensure that our results are interpretable for all and to allow comparison with earlier work, we additionally included classical statistics. Bayesian statistics allow model comparison, but also provide evidence for individual effects. When possible, we reported Bayesian model comparison (BF_10_: Bayes factor of model against the null) as well as Bayesian and frequentist effects analyses (BF_INC(LUSION)_: Bayes factor of Bayesian model averaging). We used the default JASP Cauchy priors for all Bayesian statistics^61^. Regarding frequentist statistics, we considered a p-value of 0.05 or smaller as significant. In the cases where sphericity was violated, we reported the Greenhouse-Geisser corrected p-values.

#### Color sensitivity task analysis

The data from this task were used only to establish that participants are sensitive enough to our color wheel. We calculated the overall average deviance in degrees.

#### Color wheel task data analysis

We computed the median deviance and median reaction time for all levels of ignore and update. The rationale behind choosing the median was that it is less sensitive to extreme values. For example, 90° and 180° accuracy scores are both wrong responses, but the latter affects the mean much more strongly. We used the above indices for the statistical analysis using classical and Bayesian 2×4 repeated measures ANOVAs with condition (ignore/update) and set size (levels 1-4) as within-subject factors. All participants in both experiments performed above chance level (mean deviance less than 90°).

#### Discounting choice task data analysis

As an estimate of subjective value, we computed participants’ indifference points. The indifference points can be interpreted as the financial amount offered for the presumably less effortful option (no effort or update) at which participants are equally likely to choose one or the other, thus the probability of accepting either option would be 0.5. With the main dependent variable being choice, a dichotomous variable, we calculated the probabilities of accepting the presumably less effortful offer using binomial logistic regression analysis in MATLAB and extracted the indifference points for the different conditions.

#### Choices between working memory task and taking a break

Having determined the indifference points for all levels of both working memory conditions per participant, we continued with the statistical analysis using classical and Bayesian 2×4 repeated measures ANOVAs to assess our first hypothesis that subjective value decreases with demand for ignore and update. Confirmation of this hypothesis would require that the model including set size is more likely than the null model, or the presence of a set size effect with p-value smaller than 0.05. We also performed Bayesian and classical one sample t-tests on the indifference points across levels for both conditions to assess whether the subjective value of the working memory functions was overall lower than the no task subjective value. The task offer was always €2, so a subjective value lower than 2 would imply that participants were discounting the task option.

#### Choices between Update and Ignore

We then computed participants’ indifference points collapsing across levels of “ignore vs update” choice trials to evaluate our hypothesis that ignore has a lower subjective value than update using Bayesian and classical one sample t-tests. As ignore offer was set at 2€, subjective values lower than 2 indicate that participants were willing to forgo rewards to repeat update instead of ignore trials. Additionally, we calculated indifference points for all levels separately and used a 1×4 ANOVA with set size as a factor to assess if the preference for update varies with demand.

#### Mixed effects analyses

To assess the relationship between preference and performance, we conducted various mixed effects logistic regression analyses using the lme4 package^64^ in R^65^. First, we adopted a hierarchical regression approach: we added performance as a covariate in a model regressing choice on fixed effects of set size, condition and “no effort” monetary reward. Performance scores reflected the median deviance score, per set size per condition per participant, extracted from the color wheel task. We included a random intercept per participant and by-participant random slopes for the effects of no effort monetary reward, set size and deviance. Then we ran a second, reduced model in which we removed the fixed factor condition. We compared the full with the reduced model to assess whether by adding deviance, condition no longer impacted choice. It might be noted that this model comparison approach did not allow us to include by-participant random slopes for the effect of condition, because random effects must be the same when comparing two models. For completeness, in addition to this model comparison, we ran a third model, including condition also as a by-participant random slope. In this model, we regressed choice on fixed effects of set size, condition and “no effort” monetary reward, we added a random intercept per participant and by-participant random slopes for the effects of condition and deviance. All three models were run on the ‘task vs no effort’ choice data because condition was varying by trial, in contrast to ‘ignore vs update’ choice trials, which always involved a direct comparison between conditions.

We performed similar analyses to assess whether the effect of set size can be fully accounted for by differences in performance during the color wheel task. We compared a model including set size with deviance as a covariate with a model without set size. We regressed choice on fixed factors condition, set size, deviance and monetary reward for ‘no effort’ and added a random intercept per participant with random slopes for condition, deviance and monetary reward. The optimizer used for all models was ‘bobyqa’. Model fits were compared using likelihood ratio chi-square tests.

To estimate the discounting curves across participants (Figure 3C&D) we used a mixed effects model per condition with offer amount as fixed factor and participant as a random factor.

#### Discount curve modelling

We fitted four discounting models to the “task vs no effort” choice data^23^ (Equation 1-4): linear, parabolic, hyperbolic and exponential. For this exploratory analysis, we included participants from both experiments who met the inclusion criteria for the IP analysis (74 participants). Linear models imply constant discounting with increasing effort, hyperbolic (convex) and exponential models predict greater impact of lower effort levels, while parabolic (concave) models predict greater impact of higher effort levels. We fitted the models on the ‘task versus no effort’ data per condition:

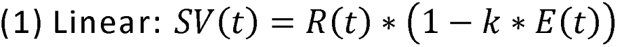

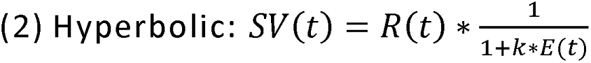

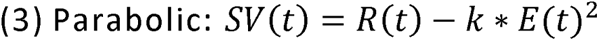

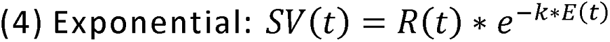

SV(t) represents the subjective value of the task option at trial t, R the magnitude of the reward offer for the no effort option, E represents the trial effort (set size) and k is the participant discounting parameter. Higher k values describe steeper discounting. The models were fit using softmax functions and maximum likelihood estimation. We compared a standard softmax function with a softmax function that includes a bias parameter^66,67^ (Equations 5 and 6) for one of the two options (effort/no effort) given the strong bias for no effort in our data.

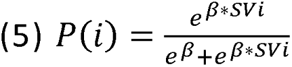

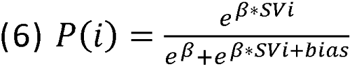

P(i) stands for the probability of choosing option i, SVi is the subjective value of option i, bias is a parameter that captures a preference for either offer irrespective of set size or reward and β is the inverse temperature of the softmax function. Each of the eight models (two softmax times four discounting functions) was fit 20 times using *fminsearch* function in Matlab, each time with different random starting points for all parameters (k, β and bias). We imposed positivity constraints on parameters k and β, while the bias parameter was left unconstrained. Model fits were compared per condition per participant with both BIC and AIC. As a predictive check, we simulated participant choices under all models using the parameters extracted per participant. Choices of the 74 participants given our task design were simulated 100 times resulting in 7400 simulated datasets per model. To compare the discounting parameters between our two conditions of interest, we conducted signed paired-sample Wilcoxon signed rank t-tests between ignore and update discounting values (k) and bias parameters. For completion, we performed the same (however unsigned) analysis for the inverse temperature of the softmax function β.

## Data availability statement

The data that support the findings of this study are available via the following persistent identifier: http://hdl.handle.net/11633/aac4rthn.

## Code availability statement

Data derivatives, analysis scripts and JASP files are provided via the following persistent identifier: http://hdl.handle.net/11633/aac4rthn.

## Author contributions

RC, DP, MIF and BZ designed the experiments, DP and MIF executed experiment 1, DP executed experiment 2, DP processed the experimental data, performed the analyses and designed the figures, DP and RC drafted the manuscript, A.W. aided in interpreting the results and offered insights in the modeling. All authors discussed the results and commented on the manuscript.

## Conflict of interest

RC and DP provide consulting services for F. Hoffmann-La Roche AG but do not hold any shares in the company.

## Acknowledgements

This research was supported by a VICI grant from NWO (Grant No. 453-14-005), an Ammodo KNAW award 2017, and a James McDonnell scholar award; all awarded to RC (220020328). We thank Rebecca Calcott^1,2^ and Lieke Hofmans^1,2^ for comments that greatly improved the manuscript, Johannes Algermissen for advice on the mixed models analyses and Trevor Chong for sharing input and scripts for the modeling analysis.

## Notes

https://github.com/danae1968/stabflex2019

### Summary of Updates

This version has been updated to better represent the cognitive processes examined, to clarify and extend some exploratory analyses and to incorporate the modeling of the discounting participant patterns.

