## Supplemental Material for "Shielding working memory from distraction is more effortful than flexible updating"

Supplementary results

| **Supplemental Table 1.** Descriptive statistics for color wheel task deviance | | | | |  |
| --- | --- | --- | --- | --- | --- |
| **Condition** | **Set size** | **Experiment 1** | | **Experiment 2** | |
|  |  | **Mean** | **SD** | **Mean** | **SD** |
| Ignore | 1 | 8.69 | 2.71 | 7.96 | 3.51 |
|  | 2 | 10.55 | 4.67 | 9.19 | 3.49 |
|  | 3 | 13.42 | 10.39 | 11.96 | 7.78 |
|  | 4 | 15.60 | 13.47 | 13.95 | 9.49 |
| Update | 1 | 8.01 | 3.24 | 7.41 | 2.84 |
|  | 2 | 7.65 | 2.70 | 8.09 | 3.76 |
|  | 3 | 8.47 | 2.60 | 8.24 | 3.51 |
|  | 4 | 10.50 | 6.02 | 11.25 | 8.94 |

| Supplemental Table 2**:** Descriptive statistics for color wheel task RTs | | | | | | |
| --- | --- | --- | --- | --- | --- | --- |
| **Condition** | **Set size** | | **Experiment 1** | | **Experiment 2** | |
|  | |  | **Mean** | **SD** | **Mean** | **SD** |
| Ignore | | 1 | 1.84 | 0.32 | 1.91 | 0.31 |
|  | | 2 | 2.27 | 0.35 | 2.25 | 0.28 |
|  | | 3 | 2.24 | 0.27 | 2.23 | 0.28 |
|  | | 4 | 2.22 | 0.29 | 2.23 | 0.30 |
| Update | | 1 | 1.93 | 0.33 | 1.94 | 0.33 |
|  | | 2 | 2.26 | 0.32 | 2.27 | 0.29 |
|  | | 3 | 2.23 | 0.31 | 2.26 | 0.26 |
|  | | 4 | 2.40 | 0.30 | 2.37 | 0.28 |

| Supplemental Table 3: Descriptive statistics for "task vs no effort" indifference points | | | | | |
| --- | --- | --- | --- | --- | --- |
| **Condition** | **Set size** | **Experiment 1** | | **Experiment 2** | |
|  |  | **Mean** | **SD** | **Mean** | **SD** |
| Ignore | Across | 1.48 | 0.45 | 1.36 | 0.52 |
|  | 1 | 1.57 | 0.47 | 1.48 | 0.49 |
|  | 2 | 1.56 | 0.39 | 1.42 | 0.52 |
|  | 3 | 1.46 | 0.48 | 1.29 | 0.58 |
|  | 4 | 1.34 | 0.56 | 1.16 | 0.62 |
| Update | Across | 1.55 | 0.46 | 1.48 | 0.42 |
|  | 1 | 1.59 | 0.44 | 1.58 | 0.41 |
|  | 2 | 1.60 | 0.44 | 1.54 | 0.45 |
|  | 3 | 1.58 | 0.50 | 1.35 | 0.55 |
|  | 4 | 1.46 | 0.57 | 1.30 | 0.56 |

| **Supplemental Table 4:** Descriptive statistics for "Ignore vs Update" indifference points | | | | |
| --- | --- | --- | --- | --- |
| **Set size** | **Experiment 1** | | **Experiment 2** | |
|  | **Mean** | **SD** | **Mean** | **SD** |
| Across | 1.88 | 0.25 | 1.73 | 0.58 |
| 1 | 1.91 | 0.22 | 1.72 | 0.42 |
| 2 | 1.88 | 0.26 | 1.71 | 0.50 |
| 3 | 1.89 | 0.24 | 1.73 | 0.69 |
| 4 | 1.85 | 0.36 | 1.73 | 0.74 |

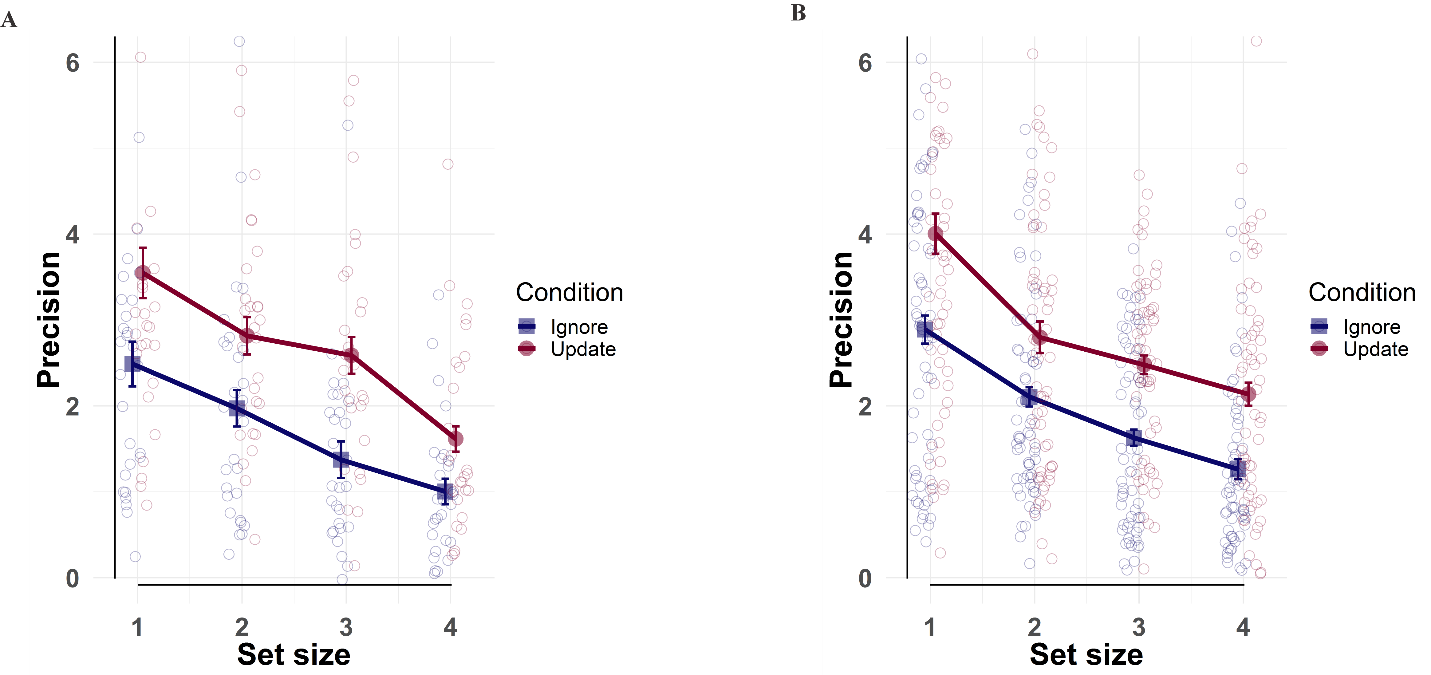
 **Supplemental** ***Figure*** ***1****.* Precision^58^ in the color wheel task as a function of set size for Experiment 1 (A, 28 participants) and Experiment 2(B, 62 participants).

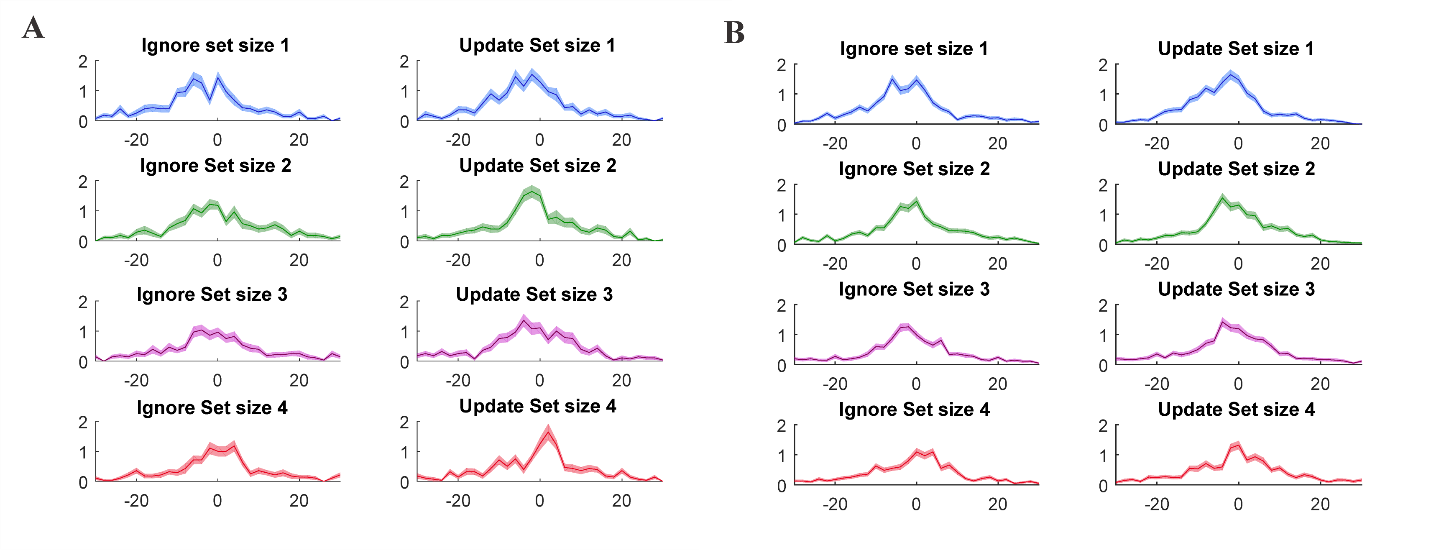

**Supplemental** ***Figure*** 2**.** Distribution of colorwheel task responses per condition per set size in Experiment 1 (A, 28 participants) and 2 (B, 61 participants). X axis represents deviance from target color.

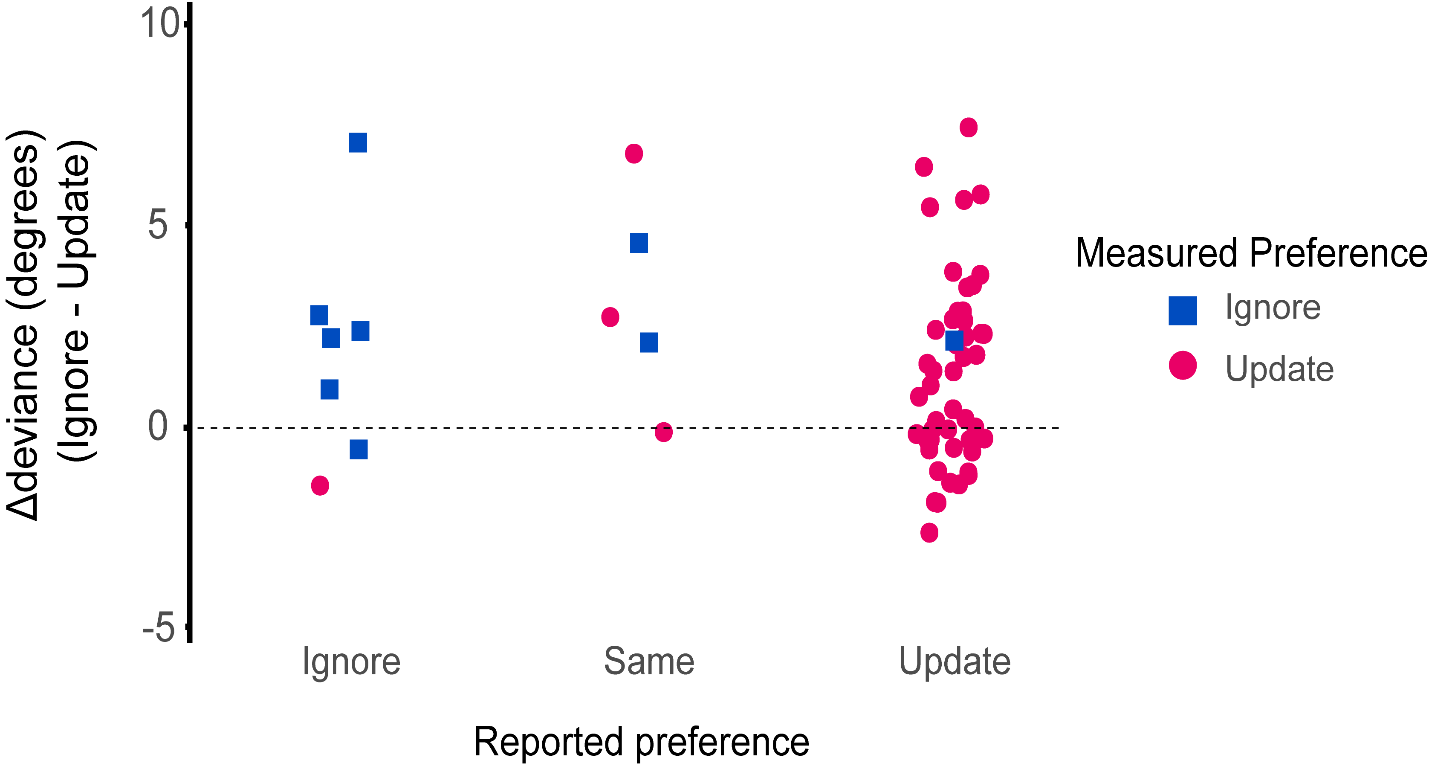

**Supplemental *Figure* 3.** Relationship between reported, measured preference and performance in the color wheel task. The y axis represents the difference in performance between ignore and update across demand. Measured preference: indifference points in the direct comparison across demand being higher or lower than 2 (preference for update or ignore respectively). Reported preference: participants’ written report of which task condition they prefer. Performance on update versus ignore trials does not covary with a preference for update versus ignore. There is a correlation between measured (indifference points) and reported (questionnaire) preference for update versus ignore. Depicted data from replication Experiment 2 (60 participants).

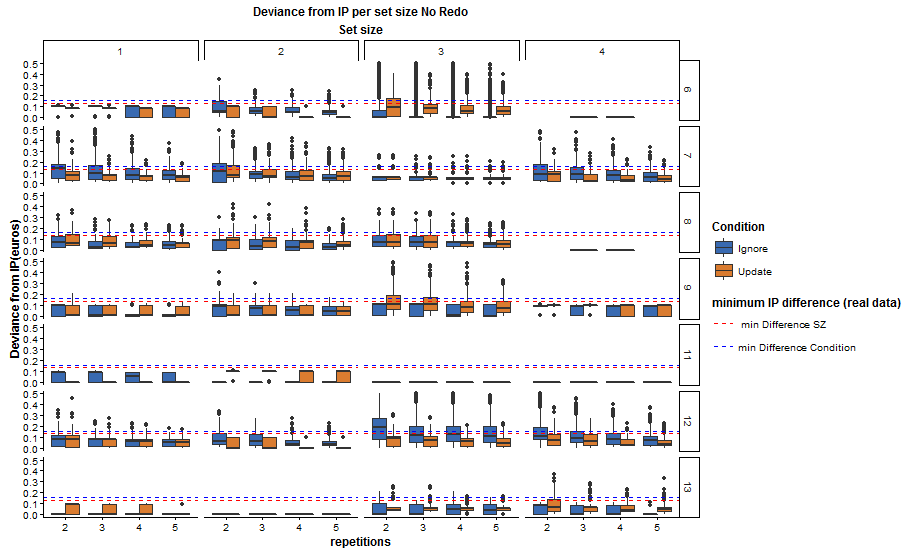
Supplemental ***Figure*** *4.* Simulations of cognitive effort discounting task given indifference points derived from pilot participants. The goal was to assess the number of repetitions of choice offer pairs necessary to extract indifference point effects for set size and condition as measured with pilot data. Every column represents a different set size, every row different participant. Every cell has been simulated 1000 times. Red line: minimum significant difference for set size derived from real data. Blue line: minimum significant condition effect difference.

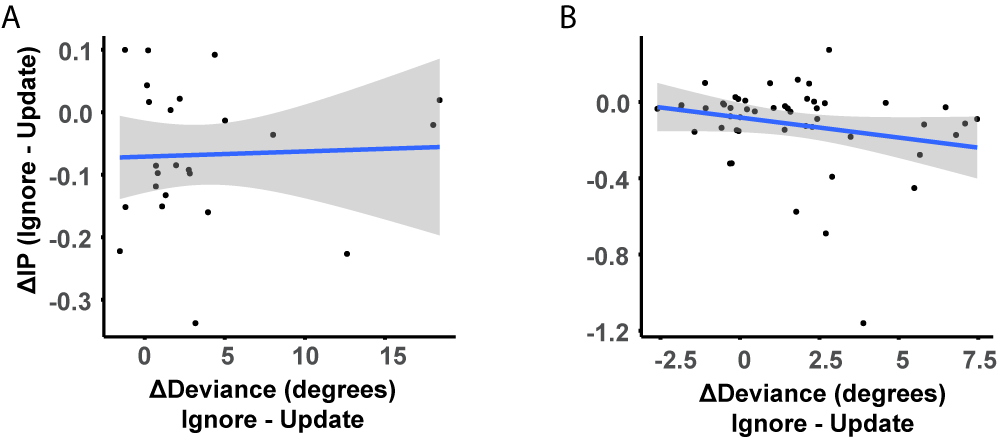

Supplemental ***Figure*** *5.* Correlation between condition differences in performance and preference (IP). Ignore minus update performance does not covary with ignore minus update IP. A Experiment 1 (Pearson’s r= 0.041, BF_10_=0.258, p=0.849, 24 participants). B Experiment 2 (Pearson’s r= -0.232, BF_10_= 0.63, p=0.106, 50 participants).

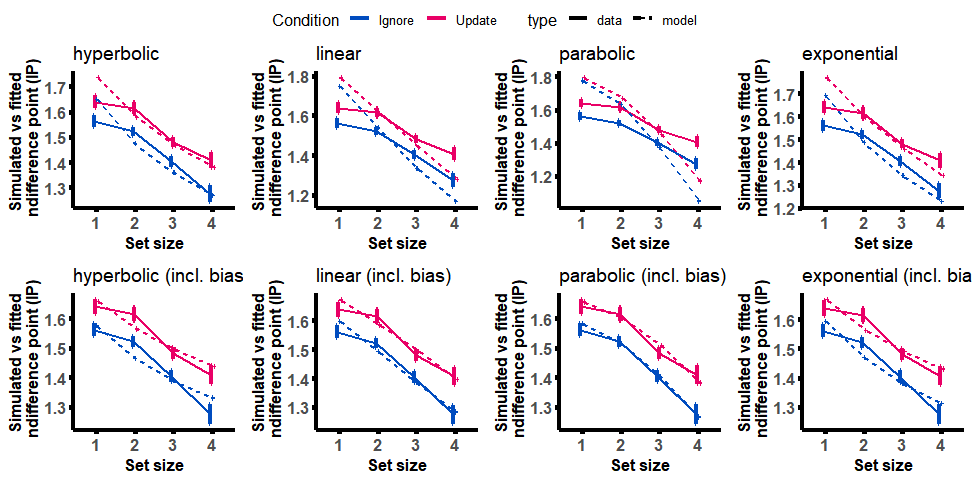
**Supplemental *Figure* 6.** Raw and simulated indifference points using the discounting parameters (k, β, bias) of all 8 fitted models (7400 simulations per model). The simulated data reproduce the effects in the observed data, even more so in the models including a bias parameter.

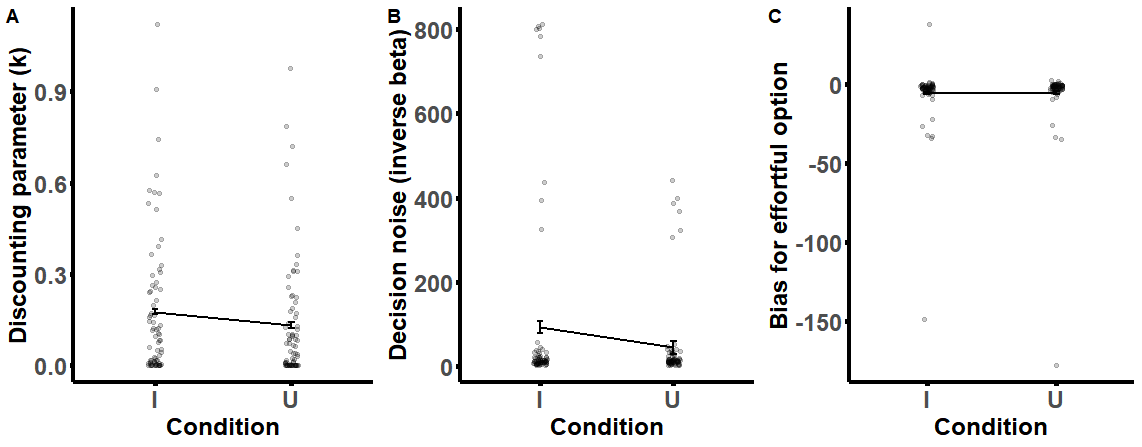

***Supplemental Figure* 7.** **A** Discounting parameter k as a function of ignore and update**. B** Decision noise β (inverse temperature of the softmax function) as a function of ignore and update. **C** Bias for/against effort option as a function of condition (n = 74, pooled across experiments).

### 1. Individual differences

In both studies, as well as in previous pilot studies, the majority of participants (83%) preferred update. However, a smaller percentage showed a preference for ignore (17%). Given this observation, we considered the possibility that, on the direct (update versus ignore) comparison trials of the COGED task, an effect of set size was in fact present, but masked by relevant individual variation in the overall preference for update or ignore. To assess this, we conducted supplementary analysis of data from both Experiment 1 and 2, in which the effect of set size on IP was stratified by a group factor representing overall preference for update over ignore. Specifically, participants were assigned to one or the other group based on their average IP being larger or smaller than 2. In keeping with our hypothesis, one-way ANOVAs in each group separately revealed effects of set size, both in those preferring update (70 participants), as well as in those preferring ignore (14 participants)(Ignore: p=0.076, BF_10_=4.56; Update: p=3.12e-4, BF_10_=865).

Supplemental Figure 8 shows that, when taking into account overall preference in the direct comparison trials, we observe an effect of set size on avoidance. This corresponds with Fig 4A&B, which illustrates data from the task-vs-no-effort trials. This effect of set size on avoidance in the direct comparison trials did not surface across the group as a whole, but only when stratifying by preference for ignore or update. In other words, those who avoided update, when averaged over all trials, were shown to avoid more difficult update trials more than easier update trials. Vice versa, those who avoided ignore, when averaged over all trials, were shown to avoid more difficult ignore trials more than lower set size ignore trials.

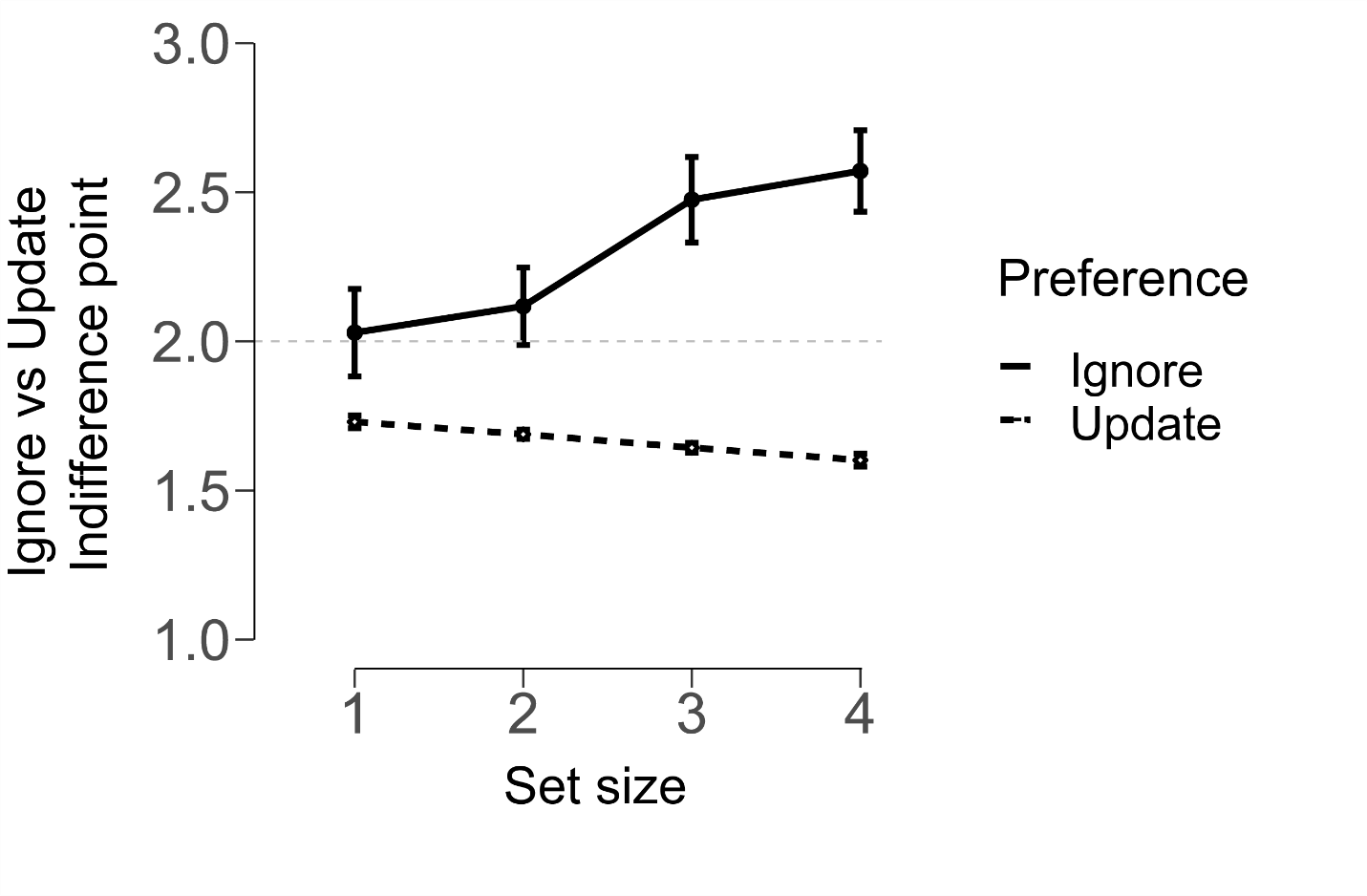

Supplemental ***Figure*** 8. “Ignore vs Update” indifference points varying by demand separately for participants who overall preferred ignore or update trials. Indifference points of 2 signify no discounting, thus, no preference between the two conditions. Participants pooled from both Experiments. The update group consists of 70 participants and the ignore group consists of 14. Error bars indicate within- participant SEM^21,22^.

### 2. Mixed effects analyses per experiment

Here, we report results of the mixed models analyses for each experiment separately. For Experiment 2, as in the case of the pooled results, the model including condition is a better fit to the data compared to the model without condition, even after including deviance (Experiment 2: model without condition: BIC: 9844, AIC: 9735; full model: BIC: 9762.9, AIC: 9646, χ^2^ = 90.9, p(pr>Chisq) < 2.2e-16). Additionally, when adding both deviance and condition as by participant random slopes the effect of condition remained significant (z = -4.77, p = 1.89e-6). For Experiment 1, we see no difference between the models (Experiment 1: model without condition: BIC: 4219.4, AIC: 4121.6; full model: BIC: 4226.9, AIC: 4122.2, χ^2^ = 1.45, p(pr>Chisq) = 0.229) and no effect of condition in model 3 (z = -1.36, (p = 0.172). However, for Experiment 1 all models gave singular fit warnings, implying that the data are not enough for this experiment to fit the complexity of the models. Moreover, when re-running the models with a different optimizer (Nelder Mead), the model including condition again outperforms the model without (model without condition: BIC: 4219.4, AIC: 4121.6; full model: BIC: 4223.4, AIC: 4118.6, χ^2^ = 4.97, p(pr>Chisq) = 0.026 Thus, the results for experiment 1 should be interpreted with caution.

### 3. Mixed effects analysis for ‘ignore vs update’ choices

Condition does not vary on every trial in the direct “Ignore vs Update” comparison trials; instead the conditions are compared directly on every trial, so we cannot assess the effect of performance on the condition effect. However, as a complimentary analysis we assess whether model fit is improved by adding performance scores in a model of direct comparison data. Performance scores reflected deviance on ignore minus update trials for the corresponding set size. We regressed choice across both experiments on the fixed effects of set size, deviance (performance score) and update monetary reward; random effects included a random intercept per participant and by-participant random slopes for the effects of update monetary reward and set size. As is the case with “No effort vs Task”, model comparison, including deviance does not improve model fit for the “Update vs Ignore” data (model without deviance: BIC: 8875, AIC: 8803; full model: BIC: 8885, AIC: 8805, p(pr > Chisq) = 0.741), converging with previous evidence that performance is not a reliable predictor of preference.
